# Hyperbolic trade-off: the importance of balancing trial and subject sample sizes in neuroimaging

**DOI:** 10.1101/2021.07.15.452548

**Authors:** Gang Chen, Daniel S. Pine, Melissa A. Brotman, Ashley R. Smith, Robert W. Cox, Paul A. Taylor, Simone P. Haller

## Abstract

Here we investigate the crucial role of trials in task-based neuroimaging from the perspectives of statistical efficiency and condition-level generalizability. Big data initiatives have gained popularity for leveraging a large sample of subjects to study a wide range of effect magnitudes in the brain. On the other hand, most taskbased FMRI designs feature a relatively small number of subjects, so that resulting parameter estimates may be associated with compromised precision. Nevertheless, little attention has been given to another important dimension of experimental design, which can equally boost a study’s statistical efficiency: the trial sample size. The common practice of condition-level modeling implicitly assumes no cross-trial variability. Here, we systematically explore the different factors that impact effect uncertainty, drawing on evidence from hierarchical modeling, simulations and an FMRI dataset of 42 subjects who completed a large number of trials of cognitive control task. We find that, due to the hyperbolic relationship between trial and subject sample sizes and the presence of relatively large cross-trial variability, 1) trial sample size has nearly the same impact as subject sample size on statistical efficiency; 2) increasing both the number of trials and subjects improves statistical efficiency more effectively than focusing on subjects alone; 3) trial sample size can be leveraged alongside subject sample size to improve the cost-effectiveness of an experimental design; 4) for small trial sample sizes, trial-level modeling, rather than condition-level modeling through summary statistics, may be necessary to accurately assess the standard error of an effect estimate. We close by making practical suggestions for improving experimental designs across neuroimaging and behavioral studies.

## 1 Introduction

Sound experimental design is key for empirical science. While reasonable statistical models may effectively extract the information of interest from the data, one first has to ensure that there is enough information present to begin with. Since there are significant constraints to acquiring data, such as cost and finite acquisition time, the experimenter should aim to optimize the experimental design to maximize relevant information within those practical limitations. A poorly designed experiment will bury signal within noise and result in unreliable findings. Of critical importance for the detection of an effect of interest in both neuroimaging and behavioral studies is to determine an appropriate sampling of a population (i.e., subjects) and a psychological process/behavior (i.e., stimuli/trials of a task/condition). Here we explore how sampling these two dimensions (i.e., subjects and trials) impacts parameter estimates and their precision. We then discuss how researchers can arrive at an efficient design given resource constraints.

### 1.1 Statistical efficiency

*Statistical efficiency* is a general metric of quality or optimization (e.g., keeping the standard error of an estimate small, while also conserving resources). One can optimize parameter estimation, a modeling framework, or an experimental design based on this quantity. A more efficient estimation process, model, or experimental design requires fewer samples than a less efficient one to achieve a common performance benchmark.

Mathematically, statistical efficiency is defined as the ratio of a sample’s inverse Fisher information to an estimator’s variance; it is dimensionless and has values between 0 and 1. However, since the model’s Fisher information is often neither known nor easily calculated, here we will refer more informally to a quantity we call “statistical efficiency” or “precision” of an effect estimate as just the inverse of the standard error. This quantity is not dimensionless and is not scaled to model information, but it conveys the relevant aspects of both the mathematical and non-technical meanings of “efficiency” for the estimation of an effect. Alternatively, we also refer to the standard error, which shares the same dimension as the underlying parameter, as a metric for the *uncertainty* about the effect estimation.

Sample size is directly associated with efficiency. As per the central limit theorem, a more efficient experimental design requires a reasonably large sample size to achieve a desired precision of effect estimation and reduce estimation uncertainty. For example, with *n* samples *x*_1_, *x*_2_,…,*x_n_* from a hypothetical population, the sample mean 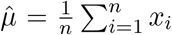 asymptotically approaches the population mean. As a study’s sample size n increases, the efficiency typically improves with an asymptotic “speed” of 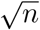 (an inverse parabola). A related concept is statistical power, which, under the conventional framework of null hypothesis significance testing, refers to the probability that the null hypothesis is correctly rejected, given that there is a “true” effect, with a certain sample size. Here, we focus on efficiency or uncertainty instead of power to broaden our discussion to a wider spectrum of modeling frameworks.

### 1.2 Subject sample size in neuroimaging

Statistical inferences are contingent on the magnitude of an effect relative to its uncertainty. For example, if the average BOLD response in a brain region is 0.8% signal change with a standard error of 0.3%, the statistical evidence is considered strong for the effect of interest. On the other hand, if the standard error is 0.6%, we would conclude that the statistical evidence for this effect is lacking because the data cannot be effectively differentiated from noise. Now if the standard error of 0.6% is based on data from only 10 participants, we may consider collecting more data before reaching the conclusion of a lack of strong evidence for the effect.

It is surprisingly difficult to predetermine an appropriate sample size in neuroimaging. In the early days a small sample size might have efficiently addressed many questions on how cognitive operations are implemented in the brain (e.g., mean brain activation in specific regions with large effect magnitudes alongside relatively low uncertainty). For example, based on data from somatosensory tasks, one study indicated that as little as 12 subjects were sufficient to detect the desired group activation patterns (without considering multiple testing adjustments) (Desmond and Glover, 2002), with 24 subjects needed to compensate for the multiplicity issue. A few power analysis methodologies have been developed over the years that are intended to assist investigators in choosing an appropriate number of subjects (e.g., fMRIPower (Mumford, 2012), Neurodesign (Durnez et al., 2016), Ostwald et al., 2019). Yet, even with these tools, power analyses are rarely performed in neuroimaging studies according to a recent survey (Szucs and Ioannidis, 2020): the median subject number was 12 among the 1,000 most cited papers during 1990-2012, and 23 among the 300 most cited papers during 2017-2018; only 3-4% of these reported pre-study power analyses. In fact, unless required for a grant application, most experiments are simply designed with sample sizes chosen to match previous studies.

Determining requisite sample sizes for neuroimaging studies is challenging. First, there is substantial heterogeneity in effect sizes across brain regions; thus, a sample size might be reasonable for some brain regions, but not for others. Second, the conventional modeling approach (massively univariate analysis followed by multiple testing adjustment) is another complicating factor. Because of the complex relationship between the strength of statistical evidence and spatial extent, it is not easy to perform power analysis while considering the multiplicity issue (e.g. permutation-based adjustment). Third, imaging analyses inherently involve multiple nested levels of data and confounds, which presents a daunting task for modeling. For instance, a typical experiment may involve several of these levels: trials, conditions (or tasks), runs, sessions, subjects, groups and population. Finally, there are also practical, non-statistical considerations involved, such as feasibility, costs, scanner availability, etc. Even though recent work has led to better characterizations of FMRI data hierarchy (Westfall et al., 2017; Chen et al., 2020; Chen et al., 2021), challenges of sample size determination remain from both a modeling and computational perspective.

Theoretically, a large subject sample size should certainly help probe effects with a small magnitude and account for a multitude of demographic, phenotypic and genetic covariates. As such, several big data initiatives have been conducted or are currently underway, including the Human Connectome Project (HCP), Adolescent Brain Cognitive Development (ABCD), Alzheimer’s Disease Neuroimaging Initiative (ADNI), Enhancing NeuroImaging Genetics through Meta Analysis (ENIGMA), UK Biobank Brain Imaging, etc. Undoubtedly, such initiatives are valuable to the research community and will continue to provide unique opportunities to explore various aspects of cognition, emotion and mental health. On the other hand, these initiatives come with high expenditure, infrastructure requirements and analytical hurdles (different sites/scanners/software). Is ‘big data’ really the best or only solution to achieving high precision for small-to-medium effects? For research questions where resources are limited (e.g., rare diseases, non-human primates), recruiting a large number of potential participants may be out of the question. In these cases, one may wonder what alternative strategies are available to achieve similar or even higher statistical efficiency with the limited number of participants or resources available.

In setting up an experiment, the choice of subject sample size is a trade-off between statistical and practical considerations. On the one hand, estimation efficiency is assumed to increase with the sample size; thus, the larger the subject sample size, the more certain the final effect estimation. On the other hand, costs (of money, time, labor etc.) increase with each added “sample” (i.e., subject); funding grants are finite, as is scanner time and even the research analyst’s time. Even though a cost-effectiveness analysis is rarely performed in practice, this trade-off does play a pivotal role for most investigations as resources are usually limited.

### 1.3 A neglected player: trial sample size

The number of trials (or data points, in resting-state or naturalistic scanning) is another important sampling dimension, yet to date it has been understudied and neglected in discussions of overall sample size. Just as the number of subjects makes up the sample size at the population level, so does the number of trials serve as the sample size for each condition or psychological process/behavior. As per probability theory’s law of large numbers, the average effect estimate for a specific condition should asymptotically approach the expected effect with increased certainty as the number of trials grows. Trial sample size often seems to be chosen for convention, practical considerations and convenience (i.e., previous studies, subject tolerance). As a result, the typical trial sample size in the field is largely in the range of [10, 40] per condition (Szucs and Ioannidis, 2020).

It seems to be a common perception that the number of trials is irrelevant to statistical efficiency at the population level, other than the need to meet a necessary minimum sample size, as evidenced by the phrase “sample size” in neuroimaging, by default, tacitly referring to the number of subjects. We hypothesize that the lack of focus on trial sample size likely results from the following two points:

- **Trial-level effects are usually of no research interest.** Often investigators are interested in condition-level effects and their comparisons. Therefore, trial-level effects generally attract little attention.
- **The conventional modeling strategy relies on condition-level summary statistics.** The conventional whole-brain, voxel-wise analysis is usually implemented in a two-step procedure: first at the subject level where trial-level effects are all bundled into one regressor (or into one set of bases) per condition; and second at the population level where cross-trial variability is invisible. Such a two-step approach avoids the computational burden of solving one “giant”, integrative model. However, as a result the cross-trial variability, as part of the hierarchical integrity of the data structure, is lost at the population level. As ultimately attention is usually paid to population-level inferences, it is this focus on condition-level effects that leads to the unawareness of the importance of both trial-level variability and trial sample size.

We note that the main goal of most FMRI studies is to generalize the results, to both the condition- and population-levels. In order to achieve these dual goals, a study must include a sufficient number of samples, in terms of *both* trials and subjects, respectively. In practice, studies tend to focus mainly on population-level level generalizability, and therefore most efforts have gone into increasing subject sample sizes (e.g., the increasing number of “big data” initiatives), while the trial sample size is typically kept at some minimal level (e.g., 20-40). As a result, we would expect the generalizability at the condition level to be challenging, in comparison to that of the population level. Condition-level generalizability is further reduced by the common modeling practice of ignoring cross-trial variability (Westfall et al., 2017; Chen et al., 2020).

A small number of studies have chosen a different strategy for experimental designs with focus on scanning subjects for an extended period of time, such as dozens of runs (e.g., Gonzalez-Castillo et al., 2012; Gordon et al., 2017), in order to obtain a large number of trials. These are variously depicted as “dense”, “deep” or “intense” sampling in the literature. Some argued that such a sampling strategy would be more advantageous due to its avoidance of potentially large cross-subject variability (Naselaris et al., 2021). Such studies should have the advantage of having high generalizability at the condition-level. However, in practice, these studies tend to include only one or a few subjects, so that generalizability to the population-level would be limited.

### 1.4 The current study

The main goal of our current investigation is to examine the impact of trial sample size (i.e., stimulus presentations) per condition alongside the number of subjects on statistical efficiency. On the one hand, the investigator does typically consider the number of trials or stimuli as a parameter during experimental design, but it is largely treated as a convenient or conventional number which the subject is able to tolerate within a scanning session. On the other hand, from the modeling perspective, the trials are usually shrouded within each condition-level regressor in the subject-level model under the assumption that all trials share exactly the same BOLD response. Furthermore, only the condition-level effect estimates are carried over to the population-level model; therefore, trial sample size does not *appear* to have much impact at the population level. However, statistically speaking the trial sample size *should* matter, because increasing the number of trials in a study increases the amount of relevant information embedded in the data. Addressing this paradox is the focus of this paper, along with the issue of study generalizability.

A related question is: *can the information associated with trial sample size be leveraged statistically to improve estimation efficiency, in the same way that increasing the number of subjects would?* It is certainly the case that increasing the number of trials in a study increases the amount of relevant information to be studied. Thus, do trial sample size and cross-trial variability play a role in statistical efficiency? And if so, how big of a role compared to the subject sample size?

In the current study, we adopt a hierarchical modeling framework, and utilize both simulations and an experimental dataset to show that trial sample size is an important dimension when optimizing task design. Importantly, we demonstrate that the “trial number” dimension has nearly the same weight and influence as its “subject number” counterpart, a fact which appears to have been underappreciated and underused in the field to date. As a result, we strongly suggest that the number of trials be leveraged alongside the number of subjects in studies, in order to more effectively achieve high statistical efficiency. In our modeling efforts, we compare the summary statistics approach of condition-level modeling (CLM) directly to a hierarchical framework of trial-level modeling (TLM) that explicitly takes cross-trial variability into consideration at the population level to examine the impact of cross-trial variability. We aim to provide a fresh perspective for experimental designs, and make a contribution to the discussion of ‘big data’ versus ‘deep scanning’ (Webb-Vargas et al., 2017; Gordon et al., 2017).

## 2 Trial-level modeling

First, we describe the formalization of our modeling framework (for convenient reference, several of the model parameters are summarized in Table 1). To frame the data generative mechanism, we adopt a simple effect structure with a group of *S* subjects who complete two task conditions (*C*_1_ and *C*_2_) while undergoing FMRI scanning. Each condition is exemplified with *T* trials (Fig. 1). We accommodate trial-level effects with a focus on the contrast between the two conditions, as is common in task FMRI. As opposed to the common practice of acquiring the condition-level effect estimates at the subject level, we obtain the trial-level effect estimates *y_cst_* of the *c*th condition (Chen et al., 2020) and assume the following effect formulation with *c, s* and *t* indexing conditions, subjects and trials, respectively:

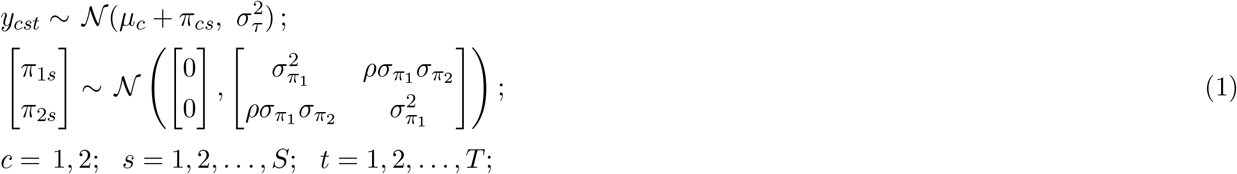

where *μ_c_* codes the population-level effect of the *c*th condition, *π_cs_* indicates the deviation of sth subject from the population effect *μ_c_* under the *c*th condition, 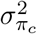 and 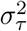 are the cross-subject and within-subject cross-trial variances, respectively, and *ρ* captures the subject-level correlation between the two conditions.

**Figure 1:**
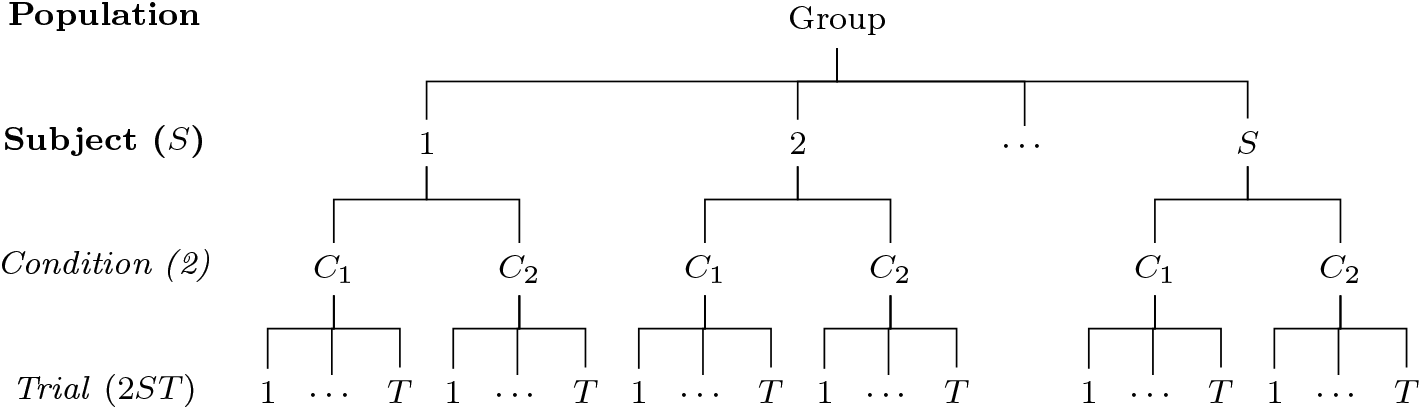
Hierarchical structure of a dataset. Assume that in a neuroimaging study a group of *S* subjects are recruited to perform a task (e.g., the Eriksen Flanker taskEriksen and Eriksen, 1974) with two conditions (e.g., congruent and incongruent) and each condition is instantiated with *T* trials. The collected data are structured across a hierarchical layout of four levels (population, subject, condition and trial) with total 2 × *S* × *T* = 2*ST* data points at the trial level compared to *S* across-condition contrasts at the subject level.

**Table 1:**
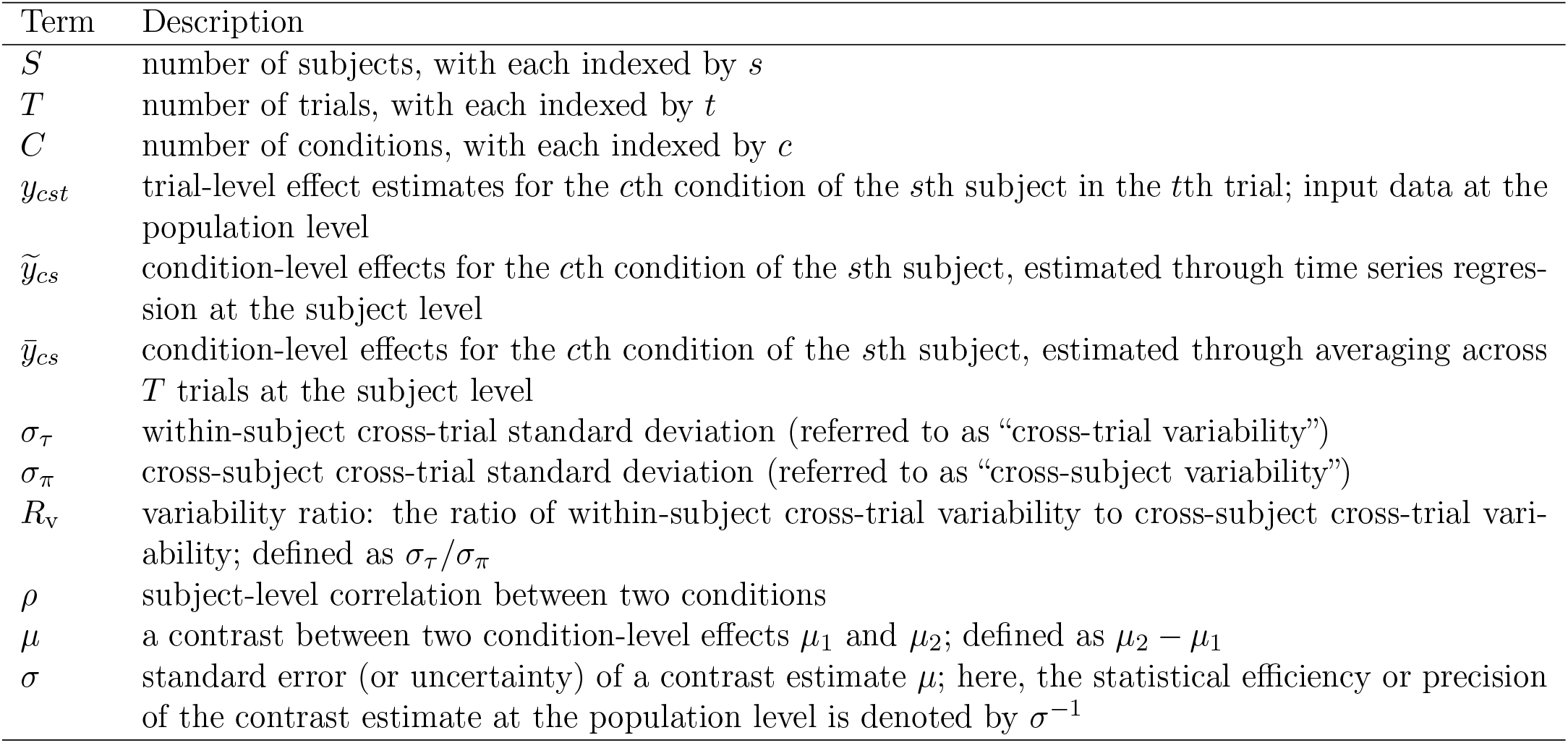
A reference table of notations used with trial-level modeling

| Term | Description |
| --- | --- |
| $S$ | number of subjects, with each indexed by $s$ |
| $T$ | number of trials, with each indexed by $t$ |
| $C$ | number of conditions, with each indexed by $c$ |
| $y_{cst}$ | trial-level effect estimates for the $c$th condition of the $s$th subject in the $t$th trial; input data at the population level |
| $\tilde{y}_{cs}$ | condition-level effects for the $c$th condition of the $s$th subject, estimated through time series regression at the subject level |
| $\bar{y}_{cs}$ | condition-level effects for the $c$th condition of the $s$th subject, estimated through averaging across $T$ trials at the subject level |
| $\sigma_\tau$ | within-subject cross-trial standard deviation (referred to as “cross-trial variability”) |
| $\sigma_\pi$ | cross-subject cross-trial standard deviation (referred to as “cross-subject variability”) |
| $R_v$ | variability ratio: the ratio of within-subject cross-trial variability to cross-subject cross-trial variability; defined as $\sigma_\tau/\sigma_\pi$ |
| $\rho$ | subject-level correlation between two conditions |
| $\mu$ | a contrast between two condition-level effects $\mu_1$ and $\mu_2$; defined as $\mu_2 - \mu_1$ |
| $\sigma$ | standard error (or uncertainty) of a contrast estimate $\mu$; here, the statistical efficiency or precision of the contrast estimate at the population level is denoted by $\sigma^{-1}$ |

One advantage of a trial-level formulation is that it allows the explicit assessment of the relative magnitude of cross-trial variability. For the convenience of discussion, we assume homoscedasticity between the two conditions: *σ*_*π*1_ = *σ*_*π*2_ = *σ*_*π*_.^1^ Specifically, the ratio of cross-trial to cross-subject variability can be defined as,

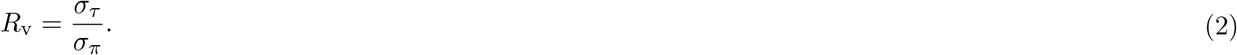

Large trial-to-trial variability has been extensively explored (He and Zempel, 2013; Trenado et al., 2019; Wolff et al., 2021). Strong evidence based on electroencephalography indicates that the substantial cross-trial variability is mainly caused by the impacts of ongoing dynamics spilling over from the prestimulus period that dwarf the influence of the trial itself (Wolff et al., 2021). Furthermore, recent investigations show that the variability ratio *R*_v_ appears often to be greater than 1 and up to 100. For example, the *R*_v_ ranged from 10 to 70 for the contrast between congruent and incongruent conditions among 12 regions in a classic Flanker FMRI experiment (Chen et al., 2021). In a reward-distraction FMRI experiment, the *R*_v_ value ranged from 5 to 80 among 11 regions (Chen et al., 2020). Even for behavioral data, which are likely significantly less noisy than neuroimaging data, the cross-trial variability is large, with *R*_v_ between 3 and 11 for reaction time data in a reward-distraction experiment (Chen et al., 2020), cognitive inhibition tasks such as the Stroop, Simon and Flanker task, digit-distance and grating orientation tasks (Rouder et al., 2019; Chen et al., 2021).

What role, if any, does trial sample size ultimately play in terms of statistical efficiency? Study descriptions typically do not discuss the reasons behind choosing their number of trials, likely a number selected by custom or convenience rather than for statistical considerations. Under the conventional analytical pipeline, each condition-level effect is estimated at the subject level through a regressor per condition. To examine differences between the conventional summary statistics pipeline through CLM and TLM as formulated in (1), we lay out the two different routes of obtaining condition-level effect estimates from subject-level analysis through time series regression: (A) obtain the *c*th condition-level effect 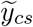 through a regressor for all the trials under the *c*th condition; (B) estimate the trial-level effects *y_cst_* using one regressor per trial and then obtain the condition-level effect through averaging,

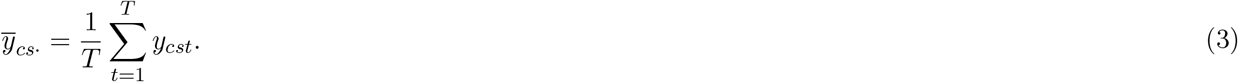

Pipeline (A) includes the following two-step process: first averaging trial-level regressors and then performing CLM through time series regression. In contrast, pipeline (B) can be considered as swapping the two steps of averaging and regression in pipeline (A): regression occurs first (i.e., TLM), followed by averaging the trial-level effect estimates. As the two processes of averaging and regression are not operationally commutative, 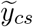 and 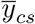 are generally not the same. However, with the assumption of an identical and independent distribution of subject-level cross-trial effects,^2^ the latter can be a proxy when we illustrate the variability of condition-level effect estimates (and later when we perform simulations of CLM in contrast to TLM):

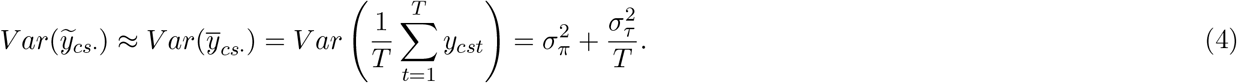

The variance expression (4) indicates that, even though trial-level effects are assumed to be the same under the conventional CLM pipeline, cross-trial variability 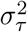 is implicitly and almost surreptitiously carried over to the population level. The important implication is that while the trial sample size *T* does not explicitly appear in the conventional CLM at the population level, it does not mean that its impact would disappear; rather, because of the way that the regressors are created, two implicit but strong assumptions are made: 1) all trials elicit exactly the same response under each condition, and 2) the condition-level effect 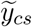 is direct measurement without any sampling error.

We now derive the expression for the standard error for the contrast between the two conditions at the population level. Directly solving the hierarchical model (1) would involve numerical iterations through, for example, restricted maximum likelihood. Fortunately, with a relatively simple data structure with two conditions, we can derive an analytic formulation that contains several illuminating features. With the notions

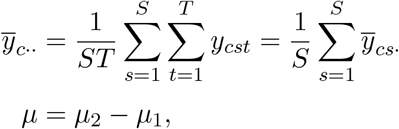

we have

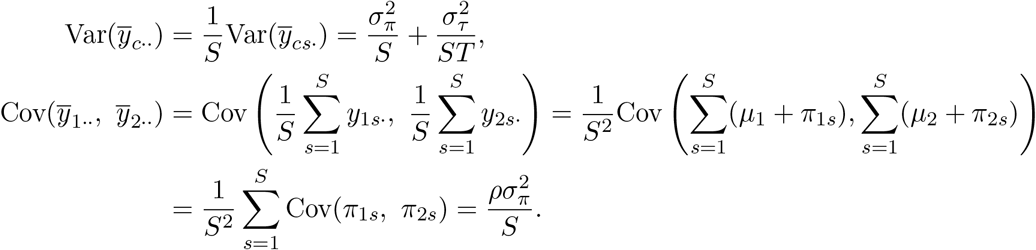

Thus, the contrast between the two conditions at the population level can be expressed as

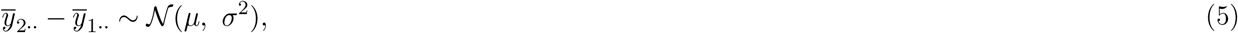

where the variance *σ*^2^ can be derived as

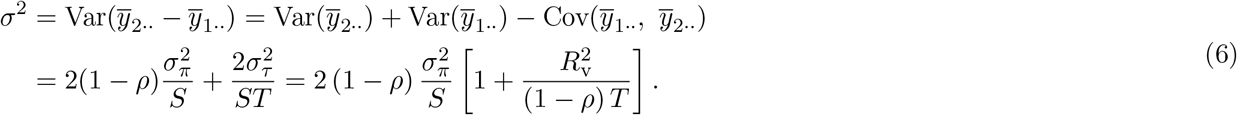

Importantly, the explicit expression for *σ*^2^ above allows us to explore the contributions of various quantities in determining the statistical efficiency for the contrast *μ*. We note that, in deriving the variance *σ*^2^, the average effects at the condition level, 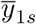 and 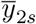, are assumed to have their respective conditional distributions; thus, trial sample size *T* and cross-trial variability *σ_τ_* directly appear in the formulation (6). In contrast, their counterparts in the conventional CLM pipeline, 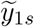 and 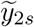, would be treated as direct measurements at the population level, leading to a one-sample (or paired) Student’s *t*-test. Below, in simulations we will use the one-sample *t*-test as an approximation for the conventional CLM pipeline and further explore this relationship. We note that it is because of this simplification in the CLM pipeline, that the impact of trial sample size *T* and cross-trial variability *σ_τ_* has been historically hidden from close examination.

The variance formula (6) has important implications for the efficiency of an experimental design or power analysis. One appealing aspect is that, when parameters *ρ, σ_π_* and *σ_τ_* are known, we might be able to find the required sample sizes *S* and *T* to achieve a designated uncertainty level *σ*. However, we face two challenges at present: the parameters *ρ, σ_π_* and *σ_τ_* are usually not empirically available; even if they were known, one cannot uniquely determine the specific sample sizes. Nevertheless, as we elaborate below, we can still gain valuable insight regarding the relationship between the subject and trial sample sizes in experimental design, as well as their impact on statistical efficiency along with the parameters *ρ, σ_π_* and *σ_τ_*.

The first variance expression (6) immediately reveals two important aspects of the two sample sizes. First, statistical efficiency, as defined as the reciprocal of the standard error *σ*, is an inverse parabolic function in terms of either the subject sample size 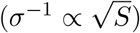 or the trial sample size 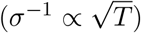. This implies that the efficiency of an experimental design improves as either sample size increases. However, this inverse parabolic relationship also means that the marginal gain of efficiency diminishes when *S* (or *T*) increases. In addition, subject sample size makes a unique contribution in the first term, which represents the cross-subject variance. The two sample sizes, *S* and *T*, combine symmetrically in the second term, which is the cross-trial variance. In the general case that the first term is not negligible compared to the second, we might say that the subject sample size influences *σ*^2^ more than the trial sample size. In the second equality of the formula (6), one sees again the direct relationship between efficiency and the cross-subject variance, which is essentially scaled by two terms: the correlation term 2 (1 – *ρ*), and the bracketed term, whose magnitude and influence is explored below.

We can rearrange the variance formula (6) and express *T* as a function of *S*, with the other quantities treated as parameters:

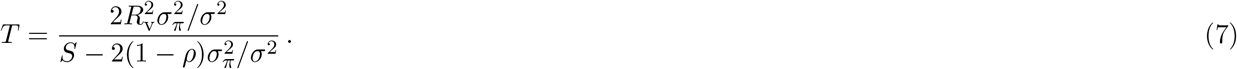

This expression shows more about the interplay between the two sample sizes within the *σ* estimation: namely that they have a hyperbolic relationship.^3^ This means that one can “trade-off” between *S* and *T* values for a given uncertainty *σ*, while all other parameters remained constant. If *σ_π_, ρ* and *R*_v_ were known, one could use the above expression to find possible combinations of *S* and *T* that are associated with a desired standard error *σ*.

Another important feature of the hyperbolic relation (7) is the presence of two asymptotes: one at *T* = *T*\* = 0, and one where the denominator is zero at

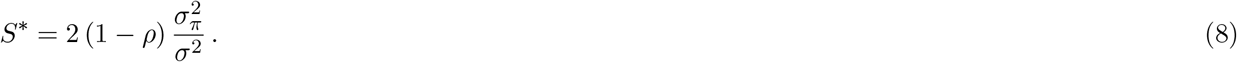

Each asymptote sets a boundary for the minimum number of respective samples required to have a given statistical efficiency (given the other parameters). For the number of trials, the requirement that *T* > *T*\* merely means there must be *some* trials acquired. For the number of subjects, *S*\* is typically nonzero, so the requirement that *S* > *S*\* can be a meaningful constraint.

These features and other relations within the expressions (6)-(7) can be appreciated with a series of example curves in Fig. 2. Each column has a fixed *ρ*, and each row has a fixed *R*_v_. Within each panel, each curve is only defined where *S* > *S*\* and *T* > *T*\*, with the vertical asymptote for each curve shown as a dotted line (and the horizontal asymptote is the *S*-axis). Each solid curve displays the set of possible (*S, T*) combinations that would result in designs having the specified *σ*, defining an isocontour of statistical efficiency. Thus, the possible trade-offs between *S* and *T* for a given *σ* are demonstrated along a curve. In terms of the “balance” of trade-offs between *S* and *T*, there are a few items to note:

1. As noted above, *S*\* sets the minimum number of subjects required to be able to reach an uncertainty level *σ*.
2. When one is near the horizontal *T* = *T*\* = 0 asymptote, there is very little marginal gain in *σ* by increasing the subject sample size *S*; this scenario corresponds to current “big data” initiatives collecting a large pool of subjects. Inversely, when approaching the vertical asymptote, we emulate the other extreme, the scenario of “deep scanning” with a lot of trials in only a few subjects, statistical efficiency barely increases when increasing the trial number. However, as indicated by the previous point, one would have to recruit a minimum number of subjects, *S*\*, to reach a designated statistical efficiency for population-level analysis (Fig. 2). In practice, the subject sample size is likely far below the threshold *S*\*.
3. Within the asymptotic region, the isocontour is symmetric around the line *T* – *T*\* = *S* – *S*\*, which simplifies here to *T* = *S* – *S*\*; that is, if (*A, B*) is a point on an isocontour, then so is (*B* + *S*\*, *A* – *S*\*).
4. Because *T*\* = 0 and *S*\* > 0, the subject sample size *S* tends to have slightly more impact on reaching a statistical efficiency than the trial sample size *T*; however, as *S*\* → 0, that difference decreases. For a given *S*\*, the amount of subject “offset” also matters less as *R*_v_ increases: the isocontour moves further from the asymptote, so the values being traded off become relatively larger, diminishing the relative impact of *S*\*. That is, in both cases, the *T* = *S* – *S*\* relation from the previous point becomes well approximated by *T* ≈ *S*, and (*S, T*) is essentially exchangeable with (*T, S*).
5. Combining the previous two points, once paying the “fixed cost” of adding the minimal number of subjects *S*\*, one can equivalently trade-off the remaining number of samples between *S* and *T*, while maintaining a constant uncertainty *σ*. Or, viewed another way, in gauging the relative importance of each sample size to reach a specific uncertainty *σ*, the number of subjects has an “extra influence” of magnitude *S*\* over the trial sample size *T*.
6. As trial number increases and *T* → ∞, the lowest uncertainty *σ* that could be achieved would be given by the first term in the variance expression (6): 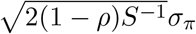.
7. The gray dashed line in Fig. 2 shows the trajectory of optimized (*S*_opt_,*T*_opt_) pairs, each defined for the constraint of having a fixed total number of samples (Appendix B). As *R*_v_ increases, the optimal trajectory approaches *S*_opt_ ≈ *T*_opt_. This is in line with the exchangeability or symmetry between the two sample sizes elaborated above.

**Figure 2:**
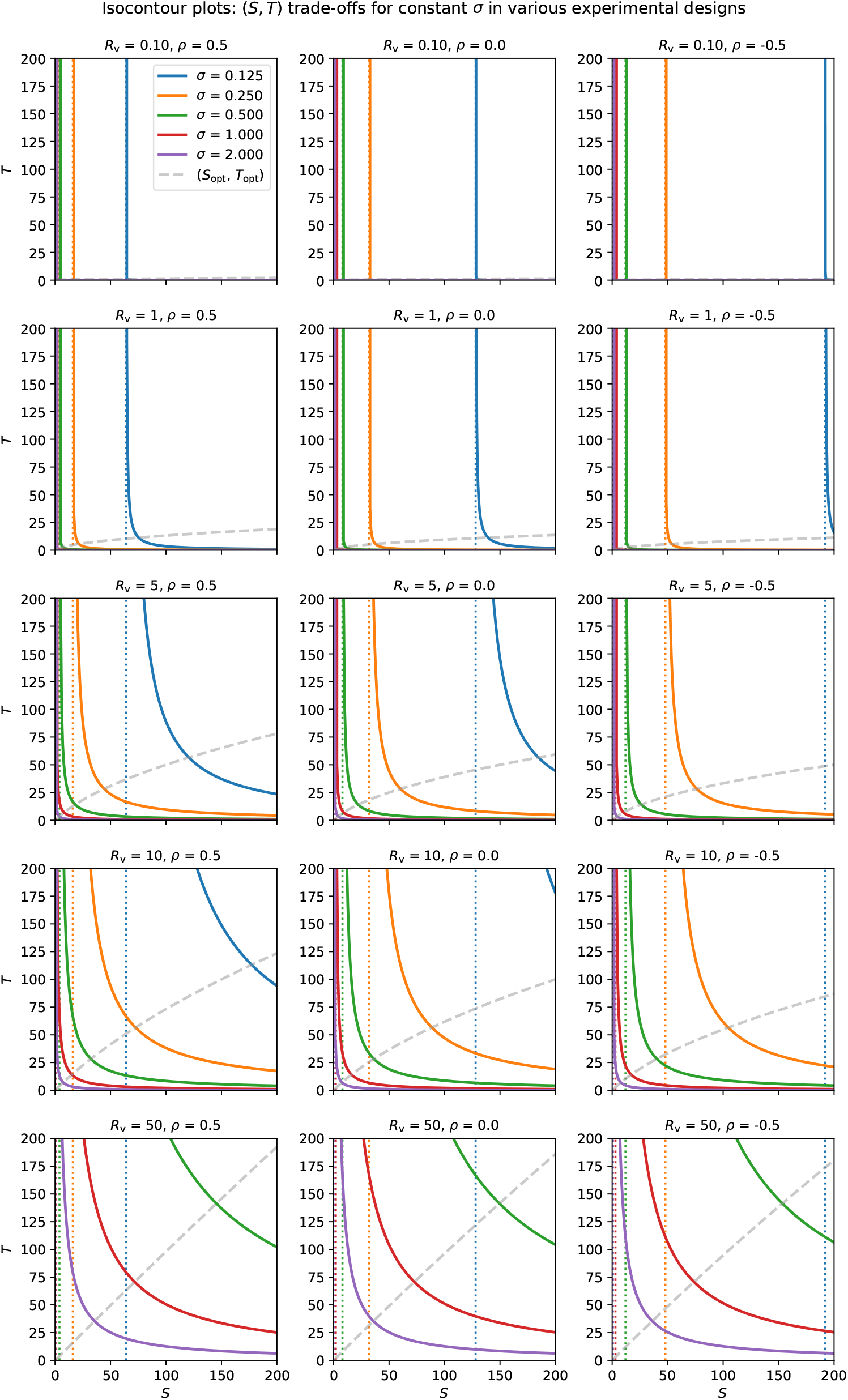
Uncertainty isocontours of subject and trial sizes. Each solid curve shows all pairs of subjects and trials that lead to the same uncertainty *σ*. The study properties are defined by the other parameters: each column shows a different value of *ρ* (0.5, 0 and −0.5), and each row has a different value of *R*_v_ (0.1, 1, 5, 10, 50). In each case, *σ*_*π*_ = 1, so *σ* has the same numerical value as *R*_v_. For a given uncertainty *σ*, there is a vertical asymptote occurring at *S*\* (dotted line, with color matching the related solid curve), which is the minimum number of subjects necessary to achieve the desired uncertainty. In the first column, the five vertical asymptotes occur (corresponding to the five *σ* values) at *S*\* = 64, 16, 4, 1, 0.25; in the second and third columns, each vertical asymptote occurs at twice and thrice the value in the first column, respectively. The gray (dashed) line shows a trajectory of (*S, T*) pairs that optimize the uncertainty *σ* for a given total number of samples (Appendix B). This (*S*_opt_,*T*_opt_) curve is nearly flat for small *R*_v_, but approaches *T* = *S* symmetry as the variability ratio *R*_v_ increases.

One can also observe from Fig. 2 the role that the correlation *ρ* plays in the estimation of *σ* and the hyperbolic relation between *S* and *T*. Namely, *ρ* does not affect the shape or slope of the hyperbola, but instead it just assists in determining the location of the *S*\* asymptote, which can be appreciated from the expressions (7)-(8). In other words, the correlation *ρ* only changes the impact of the cross-subject variance component (the first term in (7)) but not that of the cross-trial variance component (the second term in (7)). All other parameters being equal, as *ρ* increases, the minimal number of subjects *S*\* for a study design decreases. This makes intuitive sense: the smaller the correlation (including anticorrelation) between the two conditions, the more the hyperbola is shifted rightward (and thus the more difficult to detect the contrast between the two conditions). Additional views on the role of *ρ* are provided in Appendix B, in an idealized optimization case.

Finally, we note that we could express *S*\* explicitly as a function of *S* and *T* by taking the expression (8) and substituting the value of *σ*^2^ formulation (6). This results in the following relationship of *S*\* with the two sample sizes:

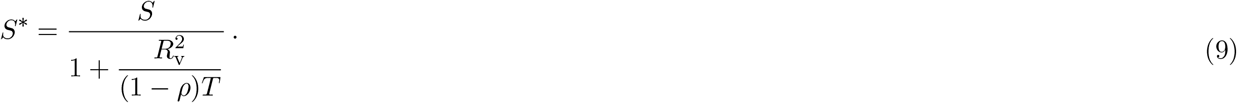

With *S*\* expressed as a function of (*S, T*), one could rearrange this expression for *T* on the left-hand side, and calculate isocontours with *S*\* held constant; these would have exactly the same shape and properties as those for uncertainty in Fig. 2. This is not surprising because there is a one-to-one correspondence between *S*\* and *σ*, as shown in the *S*\* definition (8).

### 2.1 Limit cases of trial-level modeling

To further understand the roles of various parameters, we consider different scenarios based on the variability ratio *R*_v_ and the number of trials *T*. First, we start with the second expression in the variance formulation (6), in particular the additive terms in square brackets. The second term is small when the variance ratio *R*_v_ is relatively low compared to the trial sample size *T*, producing this limiting behavior:

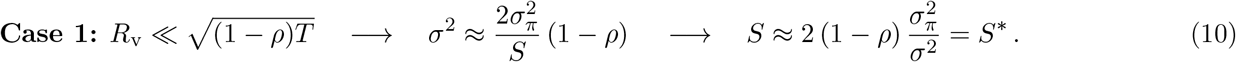

Thus, in the case of low *R*_v_ (and/or large *T*), the second component in the full variance expression could be practically ignored, and the standard error *σ* essentially depends only on the number of subjects, *σ* ∝ (*S*)^−1/2^; it is independent of the trial sample size *T* as well as cross-trial variance. For example, with 20 ≤ *T* ≤ 200 and −0.5 ≤ *ρ* ≤ 0.5, this would require that *R*_v_ be around 1 or less. In such a case, the isocontours would be approximately vertical lines, essentially matching the full contours of the first two rows in Fig. 2; and *S* is approximately the asymptotic value *S*\*. This relation also includes parts of the plots in the last three rows, as the isocontours become approximately vertical in the asymptotic limit of the trial number *T* reaching the hundreds or above.

Next, we consider the opposite limiting case. If the variability ratio *R*_v_ is relatively high compared to the trial sample size *T*, then the variance expression (6) becomes:

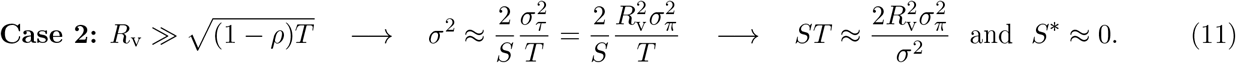

The expression for *σ*^2^ shows that standard error can be expressed independent of the cross-subject variability *σ_π_* and is dependent only on the cross-trial variability *σ_τ_*; or *σ_π_* only appears if one scales it by *R*_v_. Additionally, we note that the standard error *σ* depends on both sample sizes equally, with an asymptotic speed of *σ* ∝ (*ST*)^−1/2^. As a corollary of the relationship (9), we could say that the relative impact of *S*\* has become negligible, and so that the trade-off relationship *T* = *S* – *S*\* is well approximated by the exchange *T* ≈ *S*. Thus, the two sample sizes have equal impact on reaching an isocontour and can be equivalently traded off for each other. This is illustrated in all the isocontours except for *σ* = 0.125, 0.25 with *ρ* = 0.5 and *R*_v_ = 5 or all the isocontours except for *σ* = 0.125 (blue) with *ρ* = 0.5 and *R*_v_ = 10, 50 in Fig. 2. In practice, for typical study designs that have 20 ≤ *T* ≤ 200 and *ρ* = 0.5, this limiting case would apply if *R*_v_ were approximately greater than, for example, 20 or 100 for the respective limits.

We comment briefly on the intermediate scenario, where 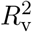 has a moderate value compared to *T*. In this case, both sample sizes play some extent of role in the uncertainty *σ*. However, as noted above, the number of subjects plays a slightly larger role than the number of trials. This is observable by the presence of a non-negligible *S*\* which offsets the (*S, T*) trade-off. In Fig. 2, relevant contours for this intermediate case are: *σ* = 0.125 (blue) with *R*_v_ = 5, 10, 50 and those of *σ* = 0.25 (blue).

We also highlight one feature of the variability ratio *R*_v_. From the above two limit cases for *σ*, we see that *R*_v_ has an important scale, based on the number of trials. That is, it is the size of *R*_v_ *relative to* 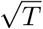 that determines much of the expected behavior of the standard error, and even whether it has any meaningful dependence on the number of trials—in Case 1, *σ* was essentially independent of *T*. The correlation *ρ* plays a role in this as well, but typically *T* is something that the experimenter controls more directly.

To summarize, we note that subject sample size *S* always plays a crucial role in achieving an adequate level of statistical efficiency. In contrast, the impact of trial sample size *T* can be much more subtle. At one extreme, its role may be negligible if *R*_v_ is around 1 or less for most trial sample sizes currently used in practice (Case 1); in fact, this limit case is what is implicitly assumed when researchers select a small number of trial repetitions and model repetitions via a single regressor. However, we emphasize that empirical data indicates that this low cross-trial variability scenario rarely occurs. At the other extreme, the trial sample size is almost as important as its subject counterpart if *R*_v_ is large relative to *T* (Case 2). In between these two limits lies the intermediate scenario where trial sample size is less important than subjects, but its influence remains sizeable. Based on the empirical values of *R*_v_, we expect that most—if not all— experimental data will likely fit into the two latter cases, with *T* being an important consideration. This has the beneficial consequence for study designs that the trial sample size can be utilized to meaningfully trade-off with the subject sample size per the variance formulation (6).

## 3 Simulations

To further explore the impact of subject and trial sizes, we use numerical simulations to test and validate the theoretical reasoning laid out above.^4^ Suppose that trial-level effects *y_cst_* are generated through the model formulation (1) with population-level effects of *μ*_1_ = 0.5, *μ*_2_ = 1.0 and a cross-subject standard deviation *σ*_π_ = 1, all in the typical BOLD units of percent signal change. Simulations for five sets of parameters were conducted:

1. five subject sample sizes: *S* = 20, 40, 60, 80, 180
2. five trial sample sizes: *T* = 20, 40, 60, 80, 180
3. five cross-trial standard deviations: *σ_τ_* = 1, 10, 20, 50, 100
4. five subject-level correlations: *ρ* = 0.1, 0.3, 0.5, 0.7, 0.9
5. two modeling approaches: trial-level (TLM) and condition-level (CLM).

With the cross-subject standard deviation set to *σ_π_* = 1, the five trial samples sizes correspond to variability ratios of *R*_v_ = 1,10,20, 50,100. With the population-level contrast *μ* = *μ*_2_ – *μ*_1_ = 0.5, the theoretical standard error is generally described by the expression (6), approaching the asymptotic expressions in (10) and (11) in some cases of *R*_v_ and 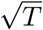. The various combinations of parameters lead to 5 × 5 × 5 × 5 × 2 = 1,250 different cases, each of which was repeated in 1000 iterations (with different random seeds).

To evaluate the simulated models we define the following quantities for investigation. For example, for model parameters such as the contrast and its standard error, we calculate the mean (or median) and standard error of each of these two estimated parameter values across 1000 iterations. Firstly, the point estimate of a parameter is considered *unbiased* when the expected mean of the sampling distribution is equal to the population value; for the present simulations, this would be the case if the mean of the estimated contrast is approximately *μ* = *μ*_2_ – *μ*_1_ = 0.5 across the iterations. Secondly, the uncertainty information for each parameter is measured in the form of 95% quantile interval. We note that the uncertainty for the effect (i.e., contrast) is numerically obtained from the repeated sampling process while the point estimate for the standard error of the effect is assessed per the respective hierarchical model. Thirdly, we validate the standard error (or efficiency) of the effect estimate per the formulation (6). Finally, we investigate the presence of a hyperbolic relationship between the two sample sizes of subjects and trials.

**Figure 3:**
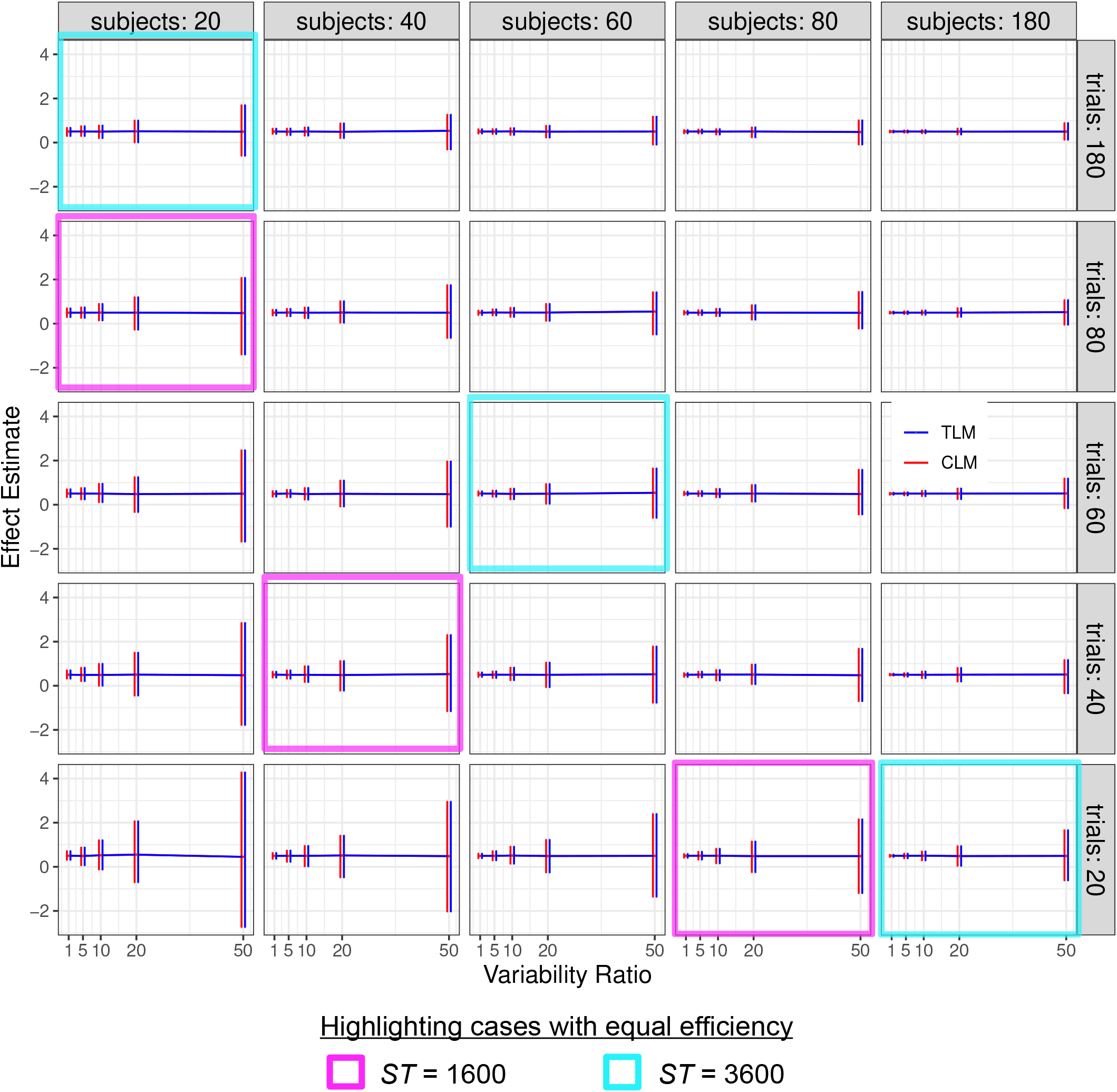
Simulation view 1: Effect estimate vs variability ratio (*x*- and *y*-axes), for various numbers of trials (panel rows) and subjects (panel columns). Results from trial-level modeling (TLM) are shown in red, and those from condition-level modeling (CLM) are shown in blue. Each horizontal line tracks the mean, and each vertical bar indicates the 95% highest density interval of effect estimates from 1000 simulations. In both cases, results typically look unbiased (the mean values are very near 0.5). Estimates are quite precise for low *R*_v_ and more uncertain as the variability ratio *R*_v_ increases, as indicated by their 95% quantile intervals. The approximate symmetry of uncertainty interval between the two sample sizes, when the variability ratio is large (e.g., *R*_v_ ≥ 10) is apparent: the magenta and cyan cells each highlight sets of simulations that have roughly equal uncertainty: note how the simulation results within each magenta block look nearly identical to each other, even though the values of *S* and *T* differ (and similarly within the cyan blocks). The correlation between the two conditions is *ρ* = 0.5; the *S, T* and *R*_v_ values are not uniformly spaced, to allow for a wider variety of behavior to be displayed.

Simulation findings are displayed in Figs. 3, 4, 5 and 6. Each plot shows a different way of “slicing” the large number of simulations, with the goal of highlighting interesting patterns and outcomes. Each plot shows results with correlations between the two conditions of *ρ* = 0.5, but the patterns and trends are quite similar for other values of *ρ*, so there is no loss of generality by simply assuming *ρ* = 0.5. As noted above, the formula (8) and Fig. 2 show that changing *ρ* typically affects the value of *S*\* and hence the location of the vertical asymptote, much more than the shape of the isocontours themselves.

**Figure 4:**
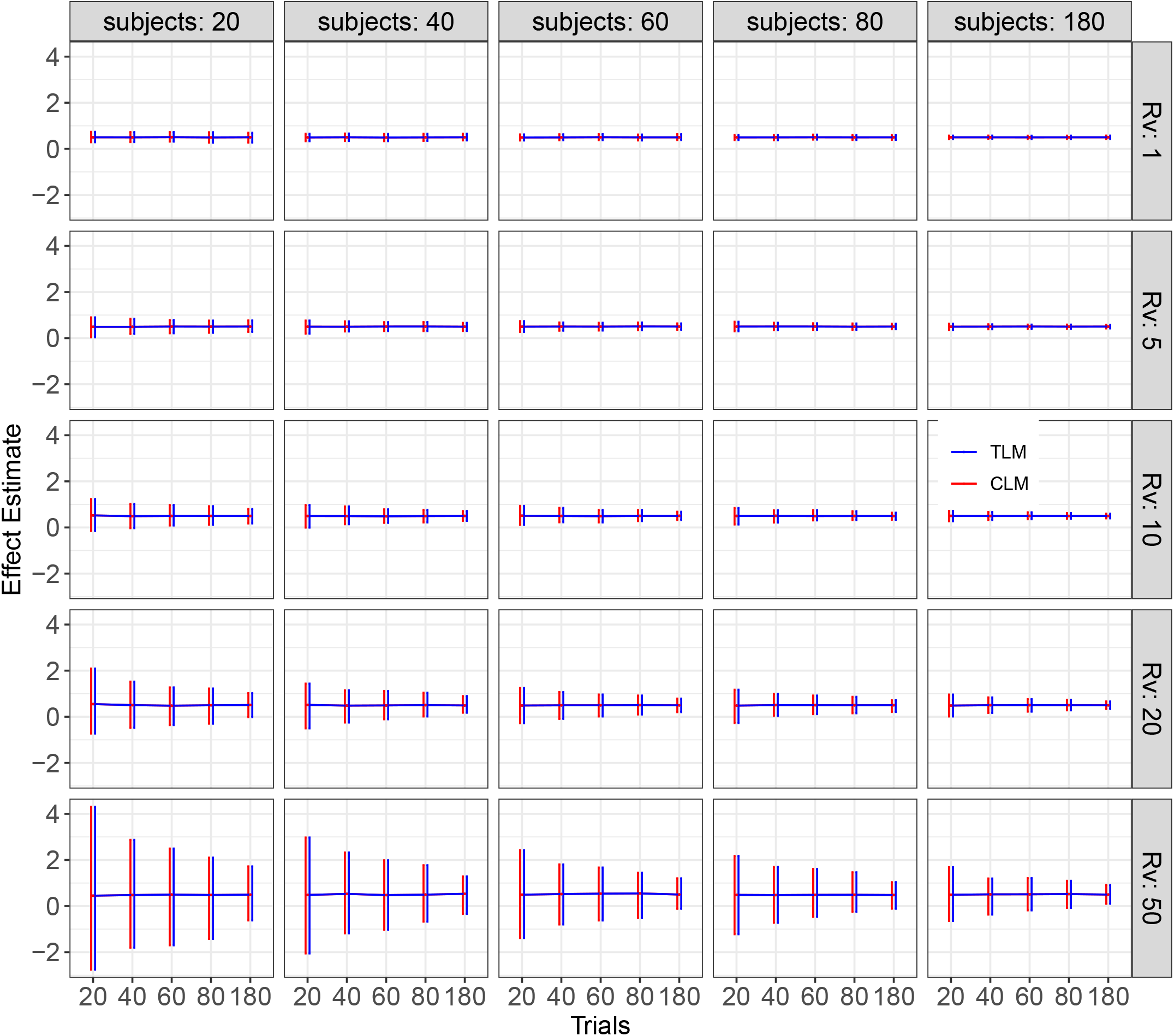
Simulation view 2: Effect estimate vs number of trials (*x*- and *y*-axes), for various variability ratios (panel rows) and numbers of subjects (panel columns). These effect estimates are the same as those shown in Fig. 3 (again, each red or blue horizontal line tracks the mean, and each bar indicates the 95% highest density interval across the 1000 simulations; *ρ* = 0.5). However, in this case the cells have been arranged to highlight the impact of the variability ratio.

**Figure 5:**
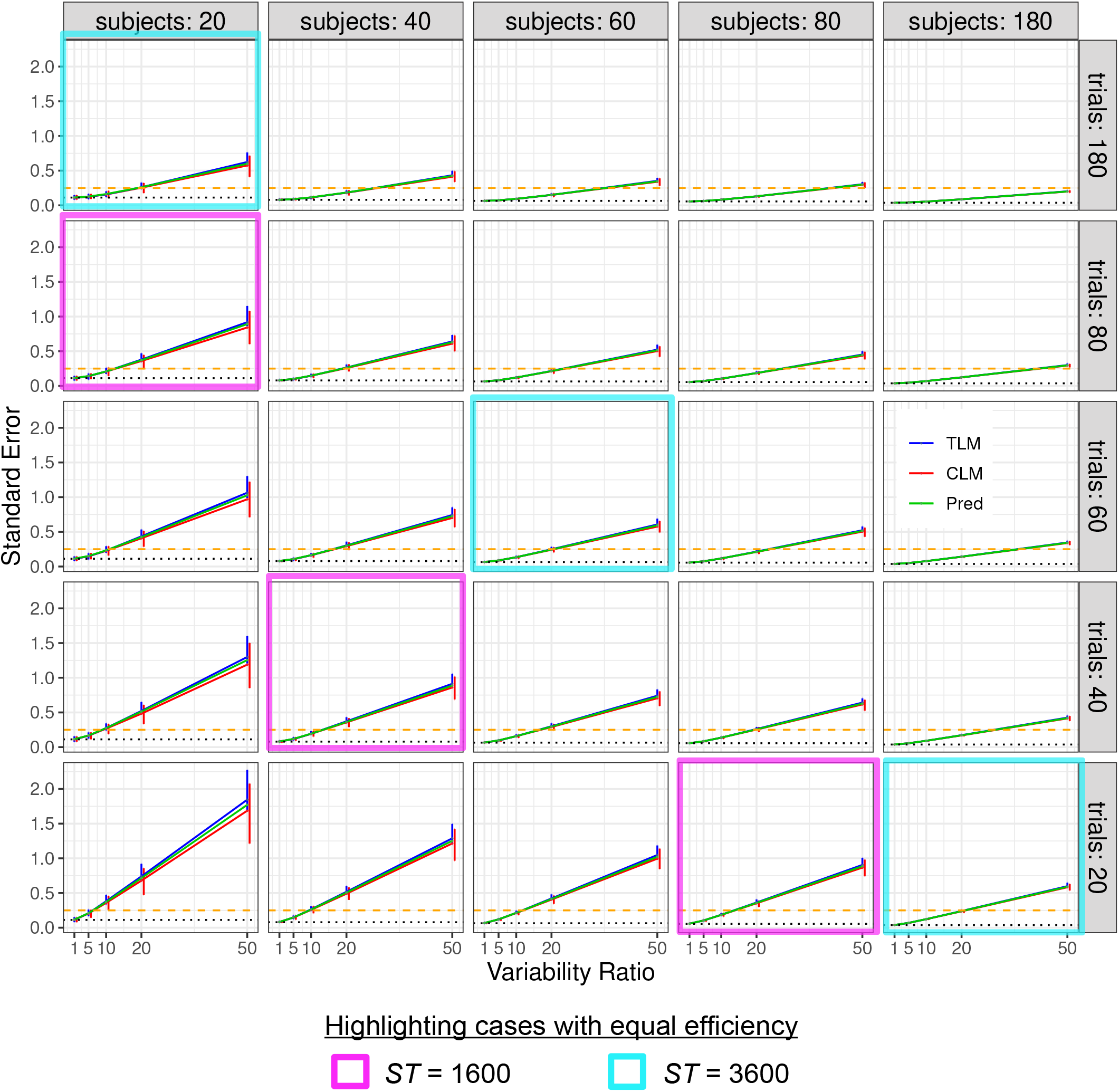
Simulation view 3: Standard error vs variability ratio (*x*- and *y*-axes), for various numbers of trials (panel rows) and subjects (panel columns). Each solid line tracks the median of the estimated standard error *σ*, and its 95% highest density interval (vertical bar) from 1000 simulations is displayed for each *R*_v_, *T* and *S*. Results from trial-level modeling (TLM) are shown in red, and those from condition-level modeling (CLM) are shown in blue; the predicted (theoretical) standard error based on the formula (6) is shown in green. The dotted line (black) marks the asymptotic standard error when the variability ratio *R*_v_ is negligible (i.e., *σ* in Case 1) or when the number of trials is infinite. The dashed line (gold) indicates the standard error of 0.25 below which the 95% quantile interval would exclude 0 with the effect magnitude of *μ* = 0.5. As in Fig. 3, one can observe the approximate symmetry between the two sample sizes when the variability ratio is large (e.g., *R*_v_ ≥ 10): the magenta and cyan cells each highlight sets of simulations that have roughly equal efficiency (cf. Fig. 3). The correlation between the two conditions is *ρ* = 0.5.

**Figure 6:**
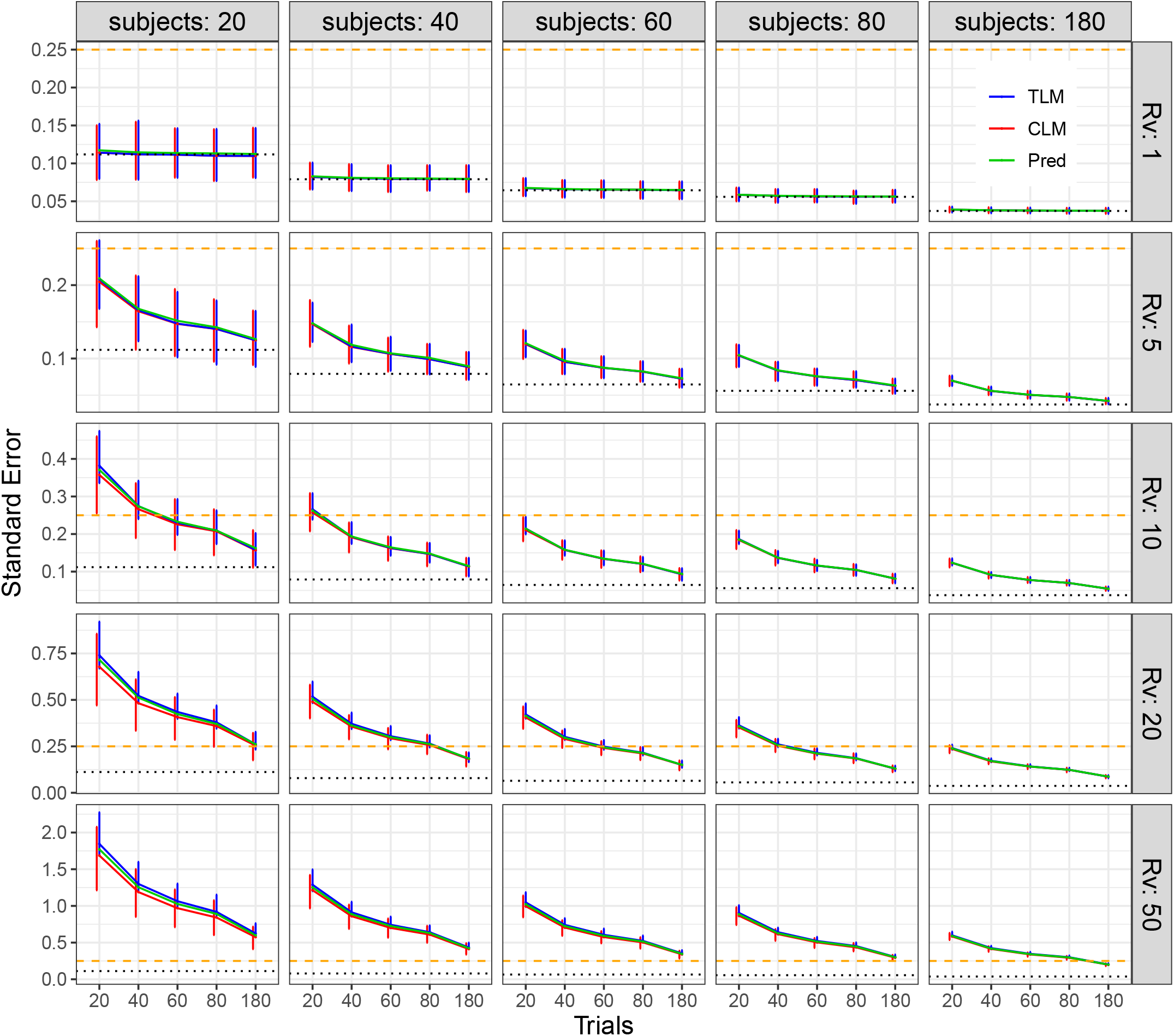
Simulation view 4: Standard error vs number of trials (*x*- and *y*-axes), for various variability ratios (panel rows) and numbers of subjects (panel columns). These standard errors are the same as those shown in Fig. 5 (again, each bar shows the 95% highest density interval across the 1000 simulations; *ρ* = 0.5). However, in this case the cells have been arranged to highlight the impact of the variability ratio, and the range of the *y*-axis in each cell varies per row. The dotted line (black) marks the asymptotic standard error when the variability ratio is negligible (i.e., *σ* in Case 1) or when the number of trials is infinite. The dashed line (gold) indicates the standard error of 0.25 below which the 95% quantile interval would exclude 0 with the effect magnitude of *μ* = 0.5.

We summarize some of the main findings from the simulations, which can be observed across the figures:

1. **Effect estimation is unbiased, but its uncertainty varies**. As shown in Figs. 3 and 4, unbiased estimation was uniformly obtained at the simulated contrast of *μ* = *μ*_2_ – *μ*_1_ = 0.5 from both TLM and CLM. However, the estimation uncertainty, as indicated by each 95% highest density interval (vertical bar) among 1000 iterations, is noticeably different across simulations. In particular, uncertainty decreases (larger bars) as *R*_v_ increases and improves (smaller bars) when the trial or subject sample size or both increase. TLM and CLM rendered virtually the same effect estimates.
2. **The standard error of effect estimation depends strongly on three factors: variability ratio, trial and subject sample sizes**. Figs. 5 and 6 show the uncertainty *σ* values from the simulations. The standard error increases with *R*_v_, and it decreases as either *T* or *S* (or *ST*) increases. Specifically, the median *σ* values from the simulations match largely well with the theoretical expectations, with TLM producing a median closer to the predictions than CLM, as well as a smaller percentile spread. The uncertainty of the effect estimate, as indicated by the 95% quantile interval (error bar) in Figs. 3 and 4, was obtained through samples across 1,000 iterations. On the other hand, the standard error estimate, another indicator of uncertainty for the effect estimate as shown in Figs. 5 and 6, was analytically assessed from the associated model. Nevertheless, these two pieces of uncertainty information for the effect estimate are comparable from each other: the 95% quantile intervals in Figs. 3 and 4 are roughly two times the standard error estimate in Figs. 5 and 6.
3. **The hyperbolic relationship is empirically confirmed in simulations.** The confirmation can be seen in the close overlap of the estimated uncertainty (blue for TLM and red for CLM in Figs. 5 and 6) versus the theoretical prediction (green). The hyperbolic relation between the number of trials and the number of subjects should allow one to trade-off *S* and *T* while keeping other parameters (e.g., statistical efficiency) constant. In addition, when the variability ratio is relatively large (e.g., *R*_v_ ≥ 10), this relationship also appeared to be validated in the symmetric pattern in these simulations—see the cyan and magenta boxes in Figs. 3 and 5, which highlight sets of cells that have roughly equal efficiency. In other words, with relatively large *R*_v_, this trade-off is directly one-for-one with an approximate symmetry (Case 2 in Sec. 2.1); for smaller variance ratios, it becomes more asymmetric and needs to trade-off a greater number of trials than subjects to keep an equal efficiency.
4. **Optimizing both trial and subject sample sizes is critical to maximize statistical efficiency.** What is an optimal way to reduce effect estimate uncertainty? Figs. 3 and 5 suggest that increasing *S* and *T* together is typically a faster way to do so than increasing either separately. For example, start in the lower left cell of Fig. 3, where *S* = *T* = 20. Note that moving to another cell vertically (increasing *T*) or horizontally (increasing *S*) leads to deceased uncertainty (smaller percentile bars). Moving either two cells up (increasing *T* by 40) or two cells right (increasing *S* by 40) leads to similar uncertainty patterns. However, moving diagonally (increasing each of *T* and *S* by 20) leads to slightly reduced uncertainty, and this property holds generally (and also for the standard error in Fig. 5). This is expected from the theoretical behavior of the hyperbolic relationship (6). In other words, these simulations reflect the fact that to maximize statistical efficiency of experimental design both sample sizes *T* and *S* should typically be increased.
5. **The differences between trial-level and condition-level modeling are subtle.** TLM and CLM rendered virtually the same effect estimates (Figs. 3 and 4). However, Figs. 5 and 6 show that CLM may result in some extent of underestimation of the standard error *σ*, as well as an increased uncertainty of *σ* (i.e., larger bars in red), in certain scenarios. The extent to which an underestimation may occur depends on three factors: *R*_v_ and the two sample sizes. Specifically, when cross-trial variability is very small (i.e., *R*_v_ ≲ 1), the underestimation of condition-level modeling is essentially negligible unless the trial sample size *T* is less than 40. On the other hand, when cross-trial variability is relatively large (i.e., *R*_v_ ≳ 20), the underestimation may become substantial especially with a small or moderate sample size (e.g., *R*_v_ = 50 with ≲ 50 subjects or trials). In addition, the underestimation is more influenced by subject sample size *S* rather than trial sample size *T*. This observation of substantial underestimation when trial sample size is not large illustrates the importance of TLM and is consistent with the recent investigations (Westfall et al., 2017; Chen et al., 2020).

We reiterate that the problems with CLM are not limited to the attenuated estimation of standard error. They are also associated with increased uncertainty across the board (larger error bars in red, Figs. 5 and 6). On the other hand, some extent of overestimation of standard error occurred for TLM under the same range of parameter values (blue lines in Figs. 5 and 6). In addition, the uncertainty of the TLM standard error estimation is right skewed (longer upper arm of each error bar in blue, Figs. 5 and 6). This was caused by a small proportion of numerical degenerative cases that were excluded from the final tallies because of algorithmic failures under the LME framework when the numerical solver got trapped at the boundary of zero standard error. Although no simple solutions to the problem are available for simulations and whole-brain voxel-level analysis, such a numerical degenerative scenario can be resolved at the region level under the Bayesian framework (Chen et al., 2020).

## 4 Assessing the impact of trial sample size in a neuroimaging dataset

### 4.1 Data description

The dataset included 42 subjects (healthy youth and adults) and was adopted from two previous studies (Smith et al., 2020; Chen et al., 2021). During FMRI scanning, subjects performed a modified Eriksen Flanker task with two trial types, congruent and incongruent: the central arrow of a vertical display pointed in either the same or opposite direction of flanking arrows, respectively (Eriksen and Eriksen, 1974). The task has a total of 432 trials for each of the two conditions, administered across 8 runs in two separate sessions. Only trials with correct responses were considered in the analysis. Thus, there were approximately 380 trials per condition per subject (350 ± 36 incongruent and 412 ± 19 congruent trials) after removing error trials.

Data processing was performed using AFNI (Cox, 1996). Details regarding image acquisition, pre-processing and subject-level analysis can be found in Appendix A. Effect estimates at the trial-level for correct responses in each condition were obtained with one regressor per trial for each subject using an autoregressive-moving-average model ARMA(1, 1) for the temporal structure of the residuals through the AFNI program 3dREMLfit (Chen et al., 2012). For comparison, effects at the condition level were also estimated through the conventional CLM approach using one regressor per condition via 3dREMLfit. The main contrast of interest was the comparison between the two conditions (i.e., incongruent versus congruent correct responses).

### 4.2 Assessing cross-trial variability across the brain

The top row of Fig. 7A displays axial slices of the effect estimate of interest, the contrast “incongruent versus congruent”. The lower row displays the variability ratio *R*_v_ associated with the contrast, which was estimated at the whole-brain voxel level through the model (1) using the AFNI program 3dLMEr (Chen et al., 2013). Translucent thresholding was applied to the overlays: results with *p* < 0.05 are opaque and outlined, and those with decreasing strength of statistical evidence are shown with increasing transparency. Substantial heterogeneity exists across the brain in terms of the relative magnitude of cross-trial variability *R*_v_ (with most high *R*_v_ ≳ 50 in low-effect regions). Fig. 7B shows the distribution of the voxelwise variability ratio *R*_v_ for the FMRI dataset, which has a mode of 20 and a 95% highest density interval [6, 86]. These *R*_v_ values are consistent with previous investigations of variability ratios in neuroimaging (Chen et al., 2021; Chen et al., 2020) and psychometric data (Rouder et al., 2019). Interestingly, many of the locations with high effect estimates (dark range and red) had relatively low variability ratios, *R*_v_ ≲ 20 (dark blue and purple colors). The regions of high contrast and strong statistical evidence (and low-medium *R*_v_) include^5^ the intraparietal sulcus area, several visual areas, premotor eye fields and inferior frontal junction, which are likely involved in the task. Most of the rest of the gray matter, as well as white matter and cerebrospinal fluid, had notably higher *R*_v_ ≥ 50.

**Figure 7:**
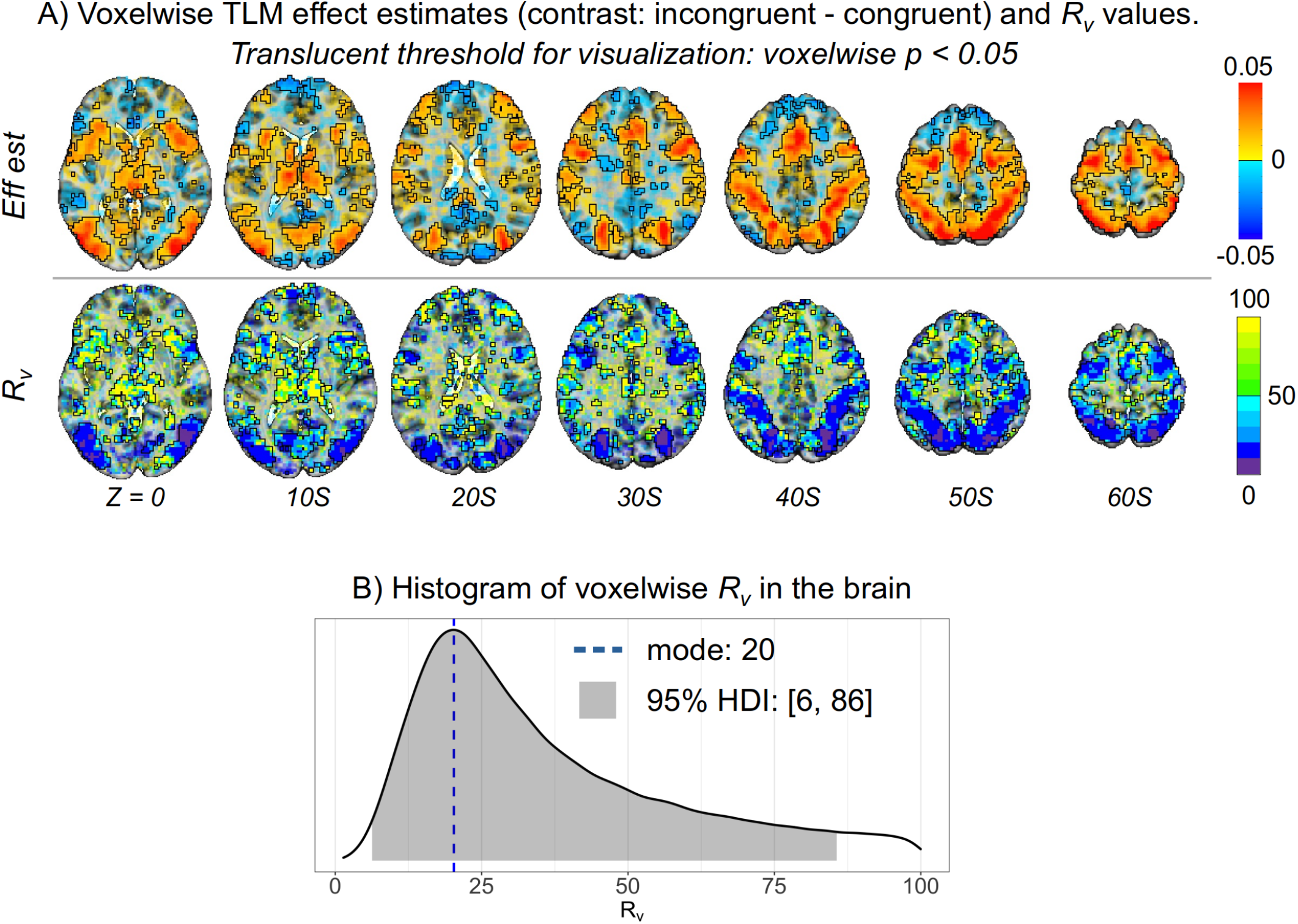
Example FMRI study, showing effect estimates and variability ratio (*R*_v_) values in the brain. The relative magnitude of cross-trial variability was estimated for the contrast “incongruent - congruent” in the Flanker dataset within the hierarchical model (1). (A) The effect estimates for the contrast and *R*_v_ values are shown in axial slices (Z coordinate in MNI standard space shown for each slice; slice orientation is in the neurological convention, right is right). For the purpose of visual clarity, a very loose voxelwise threshold of two-sided *p* < 0.05 was applied translucently: suprathreshold regions are opaque and outlined, with subthreshold voxels become increasingly transparent. Several parts of the brain have relatively low variability (*R*_v_ < 20), particularly where the contrast is largest and has strong statistical evidence. In some regions of the brain the *R*_v_ values tend to be much higher (*R*_v_ ≳ 50). (B) The mode and 95% highest density interval (HDI) for the distribution of *R*_v_ values in the brain are 20 and [6, 86], respectively.

### 4.3 Impact of trial sample size

Next, we investigated the impact of increasing trial sample size using the same Flanker FMRI dataset. Four different trial sample sizes were examined, by taking subsets of the total number available “as if” the amount of scanning had been that short: 12.5% (≈ 48 trials from the first run during the first session); 25% (≈ 95 trials from the first run of both sessions); 50% (≈ 190 trials from the first session); and 100% (≈ 380 trials). Two modeling approaches were adopted for each of the four subdatasets with different trial sample sizes: TLM through the framework (1) using the AFNI program 3dLMEr, and CLM through a paired *t*-test using the AFNI program 3dttest++.

Fig. 8 shows the values of effect estimates and the associated statistics in a representative axial slice as the number of trials increases, along with the comparisons of TLM vs CLM. Regions showing a large, positive effect (hot color locations in Fig. 8A) are fairly constant in both voxelwise value and spatial extent across trial sample sizes. Additionally, they are quite similar between the two modeling approaches, as the differences for these regions (third column) are small, particularly as the trial sample size increases. In general, regions with negative effects exhibit the most change as trial sample size increases, moving from negative values of fairly large magnitude to values of smaller magnitude; most of these regions also show corresponding weak statistical evidence (cf. Fig. 8B). The statistical evidence for regions with positive effects incrementally increases with the number of trials (Fig. 8B). The difference in statistical evidence between TLM and CLM (third column) are expressed as a ratio, centered on zero: in regions with strong statistical evidence, differences are typically small in the center of the region, with some differences at the edges; in the latter case, CLM tends to render larger values, which is consistent with having a smaller standard error *σ* for a similar effect estimate (which was observed in the simulations in Figs. 5 and 6).

**Figure 8:**
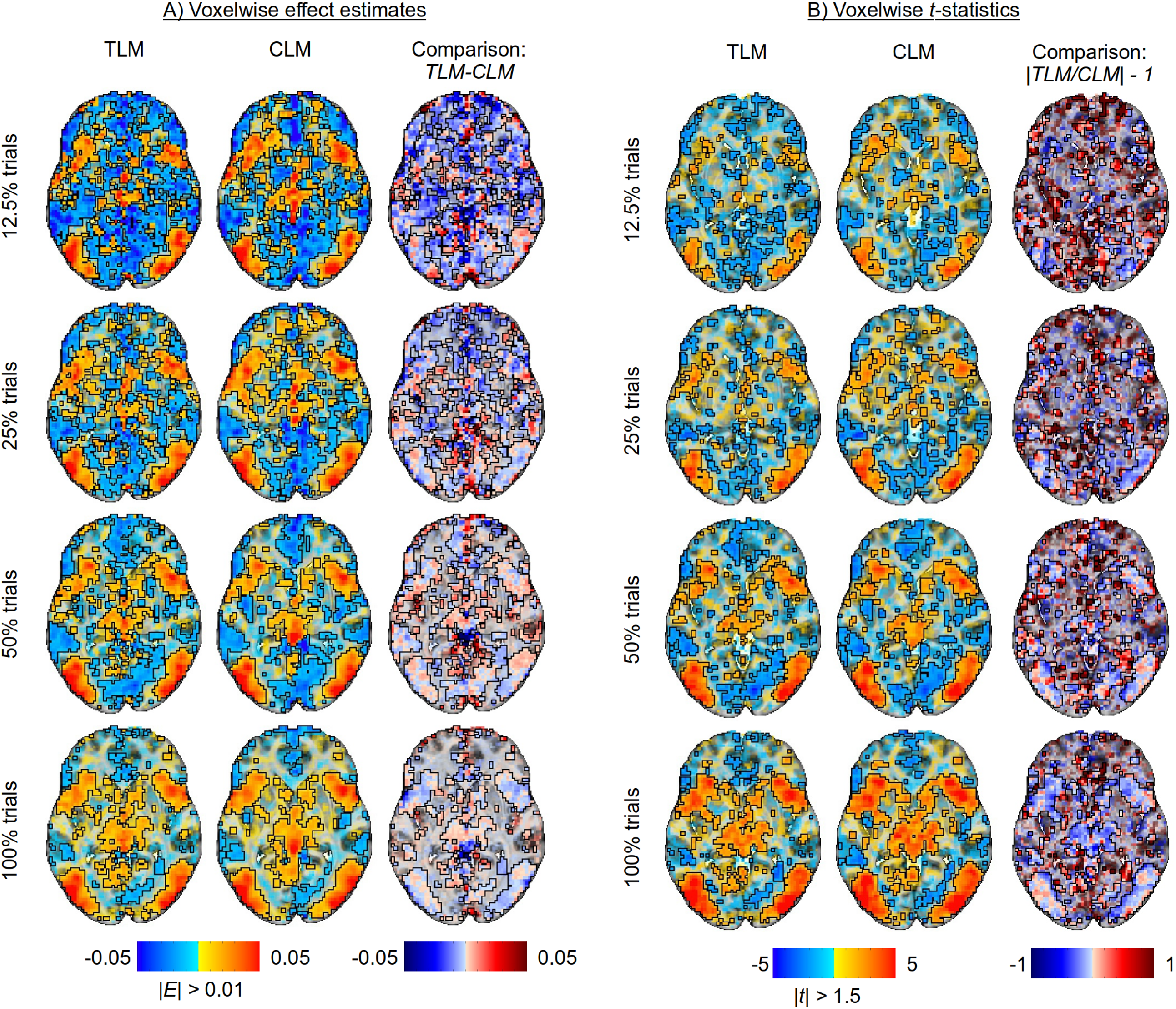
Examining differences in model outputs for both trial-level modeling (TLM) and condition-level modeling (CLM) and for various trial sample sizes (created by subsampling the full set of trials). The approximate total number of trials per subject are: 350 ± 36 incongruent trials and 412 ± 19 congruent trials. A single axial slice (*Z* = 0) is shown in each case; translucent thresholding is applied, as shown beneath the data colorbars. (A) Effect estimates of the contrast between incongurent and congruent conditions are relatively large and positive in regions with strong statistical evidence, not varying much with the number of trials or between the two modeling approaches of TLM and CLM. (B) The strength of statistical evidence for both TLM and CLM improves incrementally with the trial sample size. TLM and CLM rendered quite similar statistical results in most regions, with the latter with somewhat larger statistical values at the edges (consistent with having a similar effect estimate and smaller σ, similar to simulation results).

Fig. 9 displays a direct comparison of changes in statistical evidence with increasing number of trials. For both modeling approaches, TLM and CLM, most of the regions with positive effects show notable increase in statistical evidence with trial sample size. In scenarios where the cross-trial variability is relatively large (Case 2; cf. Fig. 7), one would theoretically expect statistical efficiency to increase with the square root of the trial sample size. Here, the number of trials doubles between two neighboring rows, which results in roughly a 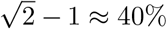 increase in the statistical value (given fairly constant effect estimates). Several parts of the plot show approximately similar rates of increase for CLM and TLM.

**Figure 9:**
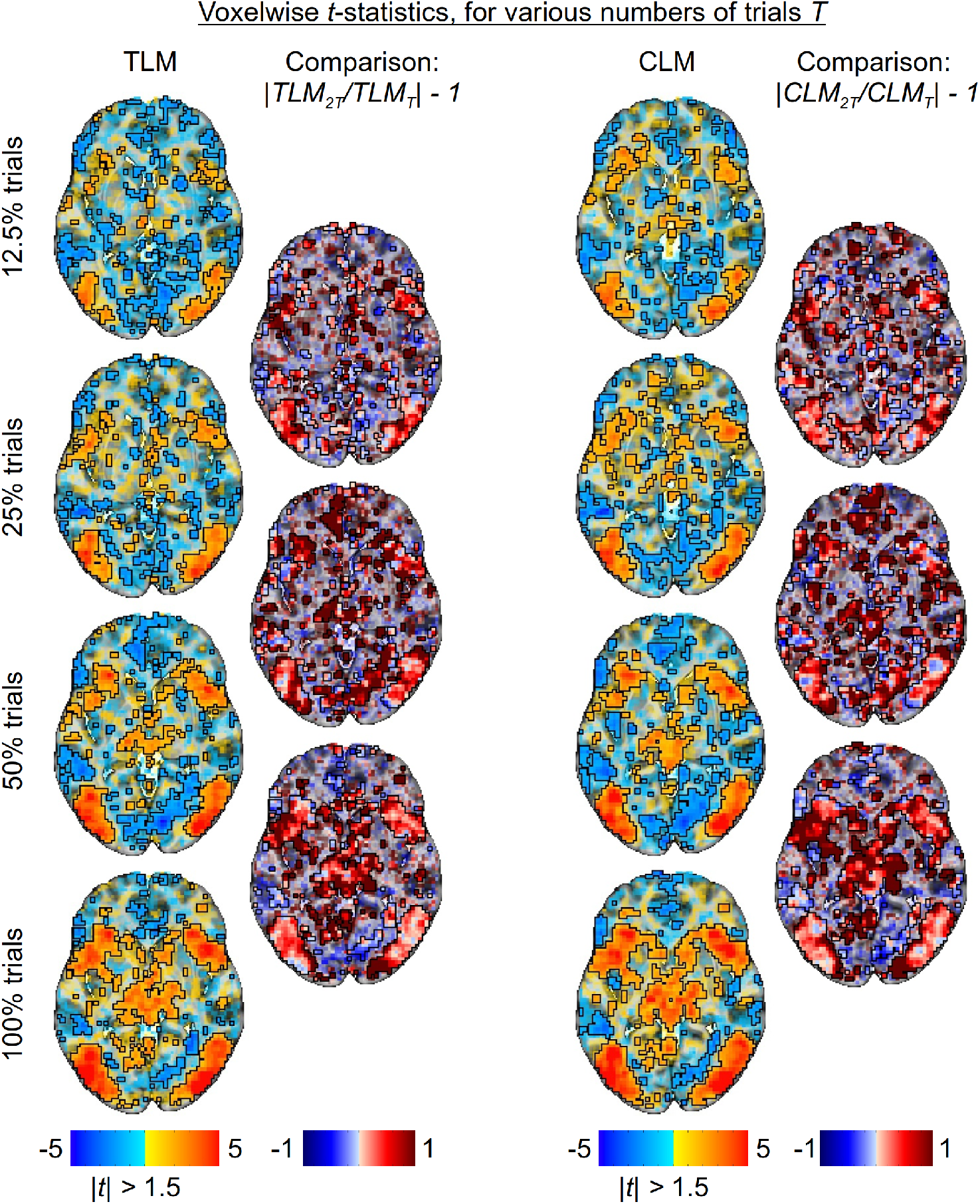
Statistical evidence with varying number of trials (*Z* = 0 axial slice). For both TLM and CLM approaches, the relative change in statistical value for the contrast between incongruent and congruent conditions, as the trial sample size doubles, is displayed as a map of the ratio of *t*-statistic magnitudes, centered on one. Thus, red shows an increase in statistical value with trial sample size and blue shows a decrease. The patterns for TLM and CLM are quite similar, increasing in most suprathreshold regions. In regions with relatively small cross-trial variability, it is expected that statistical efficiency should improve with the square of the trial sample size, since the number of trials doubles between two neighboring rows, one would expect about 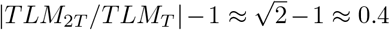 fractional increase, which is generally consistent with the results here.

## 5 Discussion

Careful experimental design is a vital component of a successful scientific investigation. An efficient design would effectively “guide” and “divert” the information to the collected data with minimal noise; an inefficient design might lead to a loss of efficiency, generalizability or applicability, or worse, a false confidence in the strength of the obtained results. Adequate sampling of both subjects and trials is crucial to detect a desired effect with adequate statistical evidence in task-related neuroimaging but also behavioral tasks. Historically, efforts to improve statistical efficiency have mostly focused on increasing subject sample size. In contrast, increasing trial number has received substantially less attention and is largely neglected in current tools to assess statistical power. In fact, there is little guidance in FMRI research on how to optimize experimental designs considering both within-subject and between-subject data acquisition. The present investigation has demonstrated how vital it is to consider trial sample size for any FMRI study.

### 5.1 Importance of trial sample size for statistical efficiency

In this investigation we show that, due to the hyperbolic relationship between the two sample sizes, the trial number plays an active role in determining statistical efficiency in a typical task-based FMRI study. Currently, many investigators tend to assume that only the number of subjects affects the efficiency of an experimental design through an inverse-parabolic relationship. Thus, most discussions of efficiency improvement focus on increasing the subject sample size, and the number of trials is largely treated as irrelevant. Assuming a negligible role for trial sample size would be valid if the cross-trial variability was small relative to cross-subject variability (first two rows in Fig. 2). However, converging evidence from empirical data indicates that the cross-trial variability is usually an order of magnitude larger than its cross-subject counterpart. As a result, trial sample size is nearly as important as subject sample size. In other words, in order to improve the efficiency of an experimental design, a large number of both trials and subjects would be optimal (see last three rows in Fig. 2). Alternatively, trial sample size can be utilized in a trade-off with the number of subjects to maintain similar design efficiency.

In practice, additional considerations such as cost, scanner time, habituation, subject fatigue, etc. will play an important role in a study design. These might affect the overall balance between the sample sizes of subjects and trials. For example, in a study of 10,000 subjects (such as the UK Biobank), it seems unfeasible to recommend having 10,000 trials per subject. Even if scan costs were covered for such a large number of trials, subject fatigue and habituation would mitigate the benefits of optimizing theoretical statistical efficiency. However, even for smaller scales in terms of number of trials, one could see efficiency benefits by having 100 versus 50 trials, for example. Hence, a positive note from the current investigation is that adding one more trial often would be more cost effective than adding one more subject; adding subjects is typically much more expensive than adding a bit more scan time. When a study includes a larger trial sample size, one might opt to tweak the design, such that multiple runs and/or multiple sessions are used to reduce fatigue. In summary, while additional practical considerations may make having roughly equal sample sizes unfeasible in some cases, most FMRI studies would benefit greatly from increasing the trial number.

The current work also sheds some light on the state of power analyses in neuroimaging. In theory, one could stick with the conventional practice in the larger field of psychology/neuroscience of estimating a study’s power (i.e., largely estimating the required sample size given a certain effect size). It is more difficult to provide an optimization tool that could assist the investigator to achieve the most efficient experimental design. Westfall et al. (2014) offered a generic power analysis interface that could aid researchers in planning studies with subject and trial sample sizes for psychological experiments. While our analyses, simulations and example experiment have shown the importance of having an adequate trial sample size, optimizing these study parameters is practically impossible given the number of unknown parameters involved in psychological and neuroimaging studies. Nevertheless, similar to trade-offs that can be made in power analyses, our discussion here emphasizes the importance of being aware of the trade-off between the two sample sizes: one can achieve the same or at least similar efficiency through manipulating the two sample sizes to a total amount of scan time or cost, or increase the efficiency by optimizing the two sample sizes with least resource cost.

### 5.2 Trial-level versus condition-level modeling: accounting for cross-trial variability

At present, most neuroimaging data analysis does not consider trial-level modeling of BOLD responses. Even in scenarios where trial-level effects are a research focus (e.g., machine learning), within-subject cross-trial variability has not been systematically investigated. Recent attempts to model FMRI task data on the trial-level (Chen et al., 2021) have revealed just how large the variance across trials within the same condition can be— namely, many times the magnitude of between-subjects variance. Traditional analysis pipelines aggregate trials into a single regressor (e.g., condition mean) per subject via condition-level modeling. This pipeline relies on the assumption that responses across all trials are exactly the same, and therefore the cross-trial variability is largely ignored. Interestingly, cross-trial variability is not even necessarily smaller for experiments that have sparse, repetitive visual displays (e.g., such as the Flanker task with simple arrow displays) compared to experiments which feature stimuli with more pronounced visual differences (e.g., smiling human faces with various “actors” who differ in their gender, age, race, and shape of facial features). Another potentially important aspect is the integration of multiple data modalities such as FMRI and EEG through hierarchical modeling (Turner et al., 2016). Similar challenges and proposals have also been discussed in psychometrics (Rouder and Haaf, 2019; Haines et al., 2020; Chen et al., 2021) as well as in other neuroimaging modalities such as PET (Chiang et al., 2017), EEG (Cai et al., 2018; Rohe et al., 2019) and MEG (Cai et al., 2018).

Cross-trial variability appears largely as random, substantial fluctuations across trials, and it is present across brain regions with no clear pattern except bilateral synchronization (Chen et al., 2020). In other words, a large proportion of cross-trial fluctuations cannot be simply explained by processes such as habituation or fatigue. However, there is some association between trial-level estimates and behavioral measures such as reaction time and stimulus ratings when modeled through trial-level modeling at the subject level. The mechanisms underlying cross-trial fluctuations remain under investigation (e.g., Wolff et al., 2021).

To more accurately characterize the data hierarchy, we advocate for explicitly accounting for cross-trial variability through trial-level modeling. Another reason for this recommendation is conceptual, researchers expect to be able to generalize from specific trials to a category or stimulus type. Simply because trial-level effects are of no interest to the investigator does not mean that they should be ignored as commonly practiced. As demonstrated in our simulations, to support valid generalizability, we suggest using trial-level modeling, especially when the trial sample size is small (i.e., 50-100 or less, Fig. 6) to avoid sizeable inflation of statistical evidence (or the underestimation of the standard error). It is worth noting that modeling at the trial-level presents some challenges at both the individual and population level. The computational cost is much higher, with a substantially larger model matrix at both the subject and population level. In addition, larger effect estimate uncertainty, outliers, and skewed distributions may occur due to high collinearity among neighboring trials or head motion; experimental design choices, such as the inter-trial interval, can be made to help reduce these issues. Recent investigations (Molloy et al. 2018; Chen et al., 2020; Chen et al., 2021) provide some solutions to handle such complex situations under the conventional and Bayesian frameworks.

### 5.3 Beyond efficiency: trial counts for generalizability, replicability, power and reliability

Properly handling uncertainty, replicability and generalizability lies at the heart of statistical inferences. The importance of considering the number of trials in a study extends beyond statistical efficiency to other prominent topics in neuroimaging. In particular, *statistical efficiency* relates to the interpretation and perception of results within a single study, but trial sample size will also have important effects on the properties that a group of studies would have—for example, if comparing results within the field or performing a meta analysis.

First, replicability within FMRI has been a recent topic of much discussion. This focuses on the consistency of results from studies that address the same research question separately, using independent data (and possibly different analysis techniques). Concerns over low rates of agreement across FMRI studies have primarily focused on the subject sample size (e.g., Turner et al., 2019). We note that replicability is essentially the same concept as uncertainty, which was discussed in Sec. 3: characterizing the spread of expected results across many iterations of Monte Carlo simulations mirrors the analysis similar datasets across groups. As shown in that section and in Figs. 5-6, increasing the number of trials plays an important role in decreasing uncertainty across iterations— and by extension, would improve replicability across studies. While some investigations of FMRI replicability have called for more data per subject (e.g., Nee, 2019), the present study provides a direct connection between the number of trials and uncertainty/replicability.

Generalizability is a related but distinct concept that refers to the validity of extending the specific research findings and conclusions from a study conducted on a particular set of samples to the population at large. Most discussions of generalizability in neuroimaging have focused on the sample size of subjects: having some minimum number of subjects to generalize to a population. However, FMRI researchers are often also interested in (tacitly, if not explicitly) generalizing across the chosen condition samples: that is, generalizing to a “population of trials” is also important. From the modeling perspective, generalizability can be characterized by the proper representation of an underlying variability through a distributional assumption. For example, under the hierarchical framework (1) for congruent and incongruent conditions, the cross-subject variability is captured by the subject-level effects *π*_1*s*_ and *π*_2*s*_ through a bivariate Gaussian distribution, while the cross-trial variability is represented by the trial-level effects through a Gaussian distribution with a variance 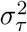. In contrast, the common practice of condition-level modeling in neuroimaging can be problematic in terms of generalizability at the condition level due to the implicit assumption of constant response across all trials (Westfall et al., 2017; Chen et al., 2020), which contradicts the reality of large cross-trial variability.

For generalizability considerations, there should also be a floor for both subject and trial sample sizes as a rule of thumb. Various recommendations have been proposed for the minimum number of subjects, ranging (at present) from 20 (Thirion et al., 2007) to 100 (Turner et al., 2018); these numbers are likely to depend strongly on experimental design, tasks, region(s) of interest (since effect magnitudes will likely vary across the brain), and other specific considerations. Similarly, no single minimum number of trials can be recommended across all studies, for the same reasons. Here, in a simple Flanker task we observed that the effect estimate fluctuated to some extent when the trial sample size changed from approx. 50 to 100 (see Fig. 8), although we note that the fluctuations were quite small in regions of strong statistical evidence. On the other hand, the same figure, along with Fig 9, shows that the statistical evidence largely kept increasing with the number of trials, even in regions that showed strong evidence at 50 trials.

Test-retest reliability can be conceptualized as another type of generalizability: namely, as the consistency of individual differences when examined as trait-like measures including behavior (e.g., RT) or BOLD response. Unlike population-level effects that are assumed to be “fixed” in a statistical model, test-retest reliability is characterized as the correlation of subject-level effects, which are termed as “random” effects under the linear mixed-effects framework. The generalizability of reliability lies in the reference of subject-level effects relative to their associated population-level effects. For example, subject-specific effects characterize the relative variations around the population effects. A high reliability of individual differences in a Flanker task experiment means that subjects with a larger cognitive conflict effect relative to the population average are expected to show a similar RT pattern when the experiment is repeated. Due to their smaller effect size compared to population effects, subject-level effects and reliability are much more subtle and may require hundreds of trials to achieve a reasonable precision for reliability estimation (Chen et al., 2021).

### 5.4 The extremes: “big data” and “deep data”

Partly to address the topics replicability and generalizability, people have proposed both *big data* studies with a large subject sample size (thousands or more) and *deep* (*or dense*) *sampling* studies with a large trial sample size (hours of scanning, several hundreds or thousands of trials). In the former, the number of trials is rarely discussed, and similarly for the latter, the number of subjects is rarely discussed. In the current context of assessing the interrelations between the two sample sizes, this means that these two options represent the extreme tails of the hyperbolic asymptotes (e.g., see Fig. 2). Additional simulation results (not shown here), which simulated these two extreme scenarios of deep scanning (3 subjects and 5000 trials) and big data (10000 subjects and 20 trials), indicated that effect estimate and uncertainty follow the general patterns summarized in Figs. 3-6: a large sample size of either trials or subjects leads to reduced standard error; a minimum number of subjects is required to achieve a designated standard error at the population level. Unsurprisingly, effect estimation for deep scanning has large uncertainty while it is relatively precise for big data.

Between these two competing opinions, big data seems to be more popular. The goals of these initiatives are to detect effects that are potentially quite small in magnitude, as well as to examine demographic variables and subgroups. However, if the number of trials is not considered as a manipulable factor in these cases, an important avenue to increased statistical efficiency is missed. In other words, an extremely large number of subjects is not necessarily the most effective way to achieve high efficiency when considering the resources and costs to recruit and collect data. Even though such large numbers of subjects would lead to the statistical efficiency gain at an asymptotic speed of inverse parabolic relationship with the number of subjects, our investigation suggests that high efficiency could be achieved with substantially fewer subjects *if* the experiment was designed to leverage the two sample sizes. Additionally, as noted above, “generalizability” comes in multiple forms, and these studies, albeit many subjects, still run the risk of not being able to properly generalize to a population of trials or stimulus category. Given the enormous cost of scanning so many subjects, this could be a lost opportunity and be inefficient, both statistically and financially. These studies might be able to save resources by scanning fewer subjects while increasing the number of trials, namely by utilizing the *S* – *T* trade-offs noted in this work. Hence, slightly smaller big data—with a larger number of trials—might be more cost-effective, similarly efficient, and generalize across more dimensions.

The other extreme of extensive sampling with a few subjects has gained attraction. For example, Gonzalez-Castillo et al. (2012) collected 500 trials per condition during 100 runs among only three subjects and revealed strong evidence to support that most brain regions are likely engaged in simple visual and attention tasks. Gordon et al. (2017) argued that a large amount of within-subject data improves precision, reliability and specificity. Naselaris et al. (2021) advocated the extensive sampling of a limited number of subjects for its higher productivity of revealing general principles. We agree that a large trial sample size with a few subjects does provide an unique opportunity to explore subject-level effects. However, we emphasize that, without an enough number of subjects to properly account for cross-subject variability, one intrinsic limitation is the lack of generalizability at the population level, as evidenced by the minimum number of subjects required for each particular uncertainty level (see *S*\* in Fig. 2). Therefore, these kinds of studies will certainly be useful for certain kinds of investigations, but the associated conclusions are usually limited and confined to those few subjects, and will not be able to generalize at the population level.

### 5.5 Limitations

We have framed study design choices and statistical efficiency under a hierarchical framework for population-level analysis. It would be useful for researchers to be able to apply this framework directly for planning the necessary numbers of subjects and trials to include in a specific study. However, in addition to effect magnitude as required in traditional power analysis, in general we do not *a priori* know the parameter values in the hierarchical model (e.g., *ρ, σ_τ_, σ_π_* in the model (1)) that would make this possible. Moreover, as seen in the Flanker dataset here, these parameter values are likely heterogeneous across the brain. In general, study designs can be much more complicated: having more conditions can lead to a more complex variance-covariance structure, for example. Some evidence indicates that the role of sample sizes may be different for classifications (such as multivoxel pattern analysis). As the spatial distribution of the effect within a region is crucial in these analytical techniques, the accuracy of statistical learning is more sensitive to cross-voxel variability than to its cross-subject and cross-trial counterparts (Davis et al., 2014).

There are limitations associated with a large number of trials. Even though increasing the number of trials can boost statistical efficiency, it must be acknowledged that this does increase scanner time and study cost. Additionally, adding trials must be done in a way that does not appreciably increase fatigue and/or habituation of the subject (particularly for youth or patient populations), otherwise the theoretical benefits to efficiency will be undermined. These practical considerations are important. Although, as noted above, in most cases adding one trial will be a noticeably lower cost than adding one subject, most studies fit scenarios where adding trials should effectively boost statistical efficiency. Splitting trials across multiple scans or sessions is one way in which this problem has been successfully approached in some large-*S* studies (e.g. **?; ?**).

The incorporation of measurement error remains a challenge within the conventional statistical framework. Specifically, FMRI acquisitions do not measure BOLD response directly, even though that is typically the effect of interest. Instead the effect is estimated from a time series regression model at the subject level. Thus, the effect estimates at the subject level, as part of the two-stage summary-statistics pipeline, contain estimation uncertainty. Ignoring this uncertainty information essentially assumes that the measurement error is either negligible or roughly the same across subjects, an assumption that simply is not met in FMRI due to noise in the acquired data and imperfect modeling. Therefore, incorporating subject-level measurement error at the population-level can calibrate and improve the model fit. Another modeling benefit is that the uncertainty information could be used to better handle outliers: instead of censoring at an artificial threshold, outliers can be down-weighting by their uncertainty (**?**). This modeling strategy has been widely adopted to achieve higher efficiency and robustness in traditional meta-analysis (**?**). Even though the methodology has been explored in straightforward population-level analyses through condition-level modeling (**?; ?; ?**), its adoption is algorithmically difficult for hierarchical modeling under the conventional statistical framework. However, it would be relatively straightforward to include the subject-level measurement errors under the Bayesian framework (**?**); yet, computational scalability presently limits its application to region-level, not whole-brain voxel-wise, analysis.

Finally, here we only examined task-based FMRI with an event-related design. It is possible that cross-trial variability and other parameters might differ for block designs. As each block with a duration of a few seconds or longer can be conceptualized as a “bundle” of many individual instantaneous trials, a block design could be viewed as more efficient than its event-related counterpart (**?**). In addition, the saturation due to the cumulative exposure, as expressed in the convolution of a block with the assumed hemodynamic response function, may also lead to much less cross-block variability compared to an event-related experiment. Resting state and naturalistic scanning are other types of popular FMRI acquisitions. Although we do not know of specific investigations examining the joint impact of number of “trials” (i.e., within-subject data) and subjects on statistical efficiency in resting-state or naturalistic scanning, we suspect that our rationale is likely applicable: the number of data points may play just as important of a role as the number of subjects in those cases (**?; ?**). Questions have been raised about the minimal number of time points needed in resting state, but not from the point of view of statistical efficiency. These are large topics requiring a separate study.

### 5.6 Suggestions and guidance

Based on our hierarchical framework, simulations and data examination, we would make the following recommendations for researchers designing and carrying out task-based FMRI studies:

1. When reporting “sample size”, researchers should be more careful and refer distinctly about “subject sample size” and “trial sample size”. Each is distinct and important in its own right.
2. When reporting the number of trials in a study, researchers should clearly note the number of trials *per condition* and *per subject*. Too often, trial counts are stated in summation, and it is not clear how many occurred per condition. This makes it difficult to parse an individual study design or for meta analyses to accurately combine multiple studies. For example, one might have to track through total trials, find the total number of participants in the final analysis, and make assumptions about relative distributions of conditions. These steps had to be performed for many entries in a recent meta analysis of highly cited FMRI studies (**?**), where a large number of papers did not even include *any* reporting of trial counts. The former situation is inexact and involves unwelcome assumptions, while the latter makes evaluating a study impossible. It should be easy for a researcher to report their trial counts per condition and per subject, to the benefit of anyone reading and interpreting the study. It would be important to provide descriptive statistics in scenarios where the final number of trials is different per subject, due to, for instance, the exclusion of trials based on subject response.
3. While we have studied the relation of statistical efficiency to subject and trial sample sizes, it is difficult to make an exact rule for choosing these sample sizes. Nevertheless, from the point of view of optimizing statistical efficiency, one could aim for a roughly equal number of trials and subjects. In practice, there are typically many more factors to consider, making a general rule difficult. However, adding more trials is *always* beneficial to statistical efficiency, and will typically improve generalizability. For example, if the resources for subjects are limited, an experiment of 50 subjects with 200 trials per condition is nearly efficient as a design of 100 subjects with 100 trials (or 500 subjects with 20 trials) per condition.
4. In addition to the statistical efficiency perspective, one should also consider generalizability, which would put a floor on both trial and subject sample sizes. It is difficult to create a single rule for either quantity, given the large variability of study designs and aims; indeed, suggestions for a minimum number of subjects have ranged from 20 to 100 (and will likely continue to fluctuate). As for trial sample size, we consider 50 as a minimum necessary number for a simple condition (e.g., the Flanker task). With a more varied task, such as displaying faces that can have a wider range of subtle but potentially important variations, using a larger number of trials would likely improve generalizability.
5. When choosing a framework between trial-level and condition-level modeling, the former is typically preferable. But the latter could be adopted when the trial sample size is reasonably large (i.e., > 50 – 100), since one might expect similar results, and condition-level modeling has the advantage of less computational burden. For smaller trial sample sizes, TLM shows clear benefits in terms of generalizability and accuracy of effect uncertainty; it is also quite computationally feasible in this range.

## 6 Conclusion

For typical neuroimaging and behavioral experiments, the trial sample size has been mostly neglected as an unimportant player in optimizing experimental designs. Large multi-site “big data” projects have proliferated in order to study small to moderate effects and individual differences. Through a hierarchical modeling framework, simulations, and an experimental dataset with a large number of trials, we hope that our investigation of the intricate relationship between the subject and trial sample sizes has illustrated the pivotal role of trials in designing a statistically efficient study. With the recent discovery that cross-trial variability is an order of magnitude higher than between-subject variability, a statistically efficient design would employ the balance of both trials and subjects. Additional practical factors such as subject tolerance and cost/resources would also need to be considered, but the importance of trial sample size is demonstrated herein.

## 7 Acknowledgments

The research and writing of the paper were supported (GC, PAT and RWC) by the NIMH and NINDS Intramural Research Programs (ZICMH002888) of the NIH/HHS, USA. Data collection was supported (DSP) by the NIMH Intramural Research Program (ZIAMH002781). This work utilized the computational resources of the NIH HPC Biowulf cluster (https://hpc.nih.gov).

## Appendices

### A Flanker Image acquisition and preprocessing

#### Image Acquisition

All procedures were approved by the National Institute of Mental Health Institutional Review Board. Participants provided written informed consent; for youth, parents provided written informed consent, while youth provided assent. Neuroimaging data were acquired from 24 adults (> 18 years; age: 26.81 ± 6.36 years) and 18 youth (< 18 years; age: 14.01 ± 2.48 years) on a 3T GE Scanner using a 32-channel head coil across two separate sessions. After a sagittal localizer scan, an automated shim calibrated the magnetic field to decrease signal dropout from a susceptibility artifact. Echoplanar images were acquired during two sessions at the following specifications: flip angle = 60°, echo time = 25 ms, repetition time = 2000 ms, 170 volumes per run, four runs per session, with an acquisition voxel size of 2.5 × 2.5 × 3 mm. The first 4 volumes from each run were discarded during pre-processing to ensure that longitudinal magnetization equilibrium was reached. Structural images were collected using a high-resolution T1-weighted magnetization-prepared rapid acquisition gradient echo (MPRAGE) sequence for co-registration with the functional data. Images were collected with a flip angle of 7° at a voxel size of 1 mm isotropic.

#### Image Pre-processing

Neuroimaging data were processed and checked using AFNI version 20.3.00 (**?**). Standard single subject pre-processing included specifying the following with afni_proc.py: despiking, slicetiming correction, distortion correction, alignment of all volumes to a base volume with minimum outliers, nonlinear registration to the MNI template, spatial smoothing to a 6.5 mm FWHM kernel, masking, and intensity scaling. Final voxel size was 2.5 × 2.5 × 2.5 mm. We excluded any pair of successive TRs in which the sum head displacement (Euclidean norm of the derivative of the translation and rotation parameters) between those TRs exceeded 1 mm. TRs in which more than 10% of voxels were outliers were also excluded. Participants’ datasets were excluded if the average motion per TR after censoring was greater than 0.25 mm or if more than 15% of TRs were censored for motion or outliers. In addition, 6 head motion parameters were included as nuisance regressors in individual-level models.

#### Subject-level Analysis

At the subject level, we analyzed brain activity with a time series model with regressors time-locked to stimulus onset reflecting trial type (incongruent, congruent), also using afni_proc.py. Regressors were created with a gamma variate for the hemodynamic response. The effect of interest at the condition level was the cognitive conflict contrast (“incongruent correct responses” versus “congruent correct responses”) with a total of 32,005 observations across two sessions of the Flanker task, which corresponds to approximately 350±36 incongruent trials and 412± 19 congruent trials per subject. We compare two approaches at the whole-brain level: a conventional condition-level modeling with regressors created at the condition level and trial-level modeling with trial-level regressors.

### B Determining optimal sample sizes

The question of how to optimize the selection of subject sample size *S* and trial sample size *T* in an experiment, given the constraint of a fixed total number of samples, can be addressed with the classical method of Lagrange multipliers. Let **X** be a 1D vector of variables, *f*(**X**) be the objective function whose extrema are to be found, and *g*(**X**) = 0 express the constraint. Then we can create the Lagrange function:

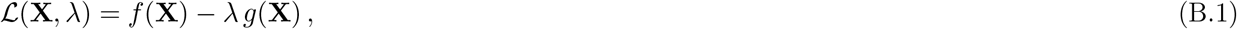

where λ is an unknown scalar parameter, introduced temporarily but not affecting final values.

For the present study, we have **X** = (*S, T*), and *N* = *S* + *T* represents the total number of samples to be partitioned between subjects and trials. We want to find the values of (*S, T*) for which *σ*^2^ is a minimum (i.e., minimal uncertainty or maximal efficiency) for an experimental design constrained to have *N* total samples, and for which *R*_v_, *σ_π_, σ_τ_* and *ρ* are just constant parameters. Therefore, we use the variance expression (6) as the objective function *f*(*S, T*). The formula for the number of total samples *N* can be fitted into the following constraint expression,

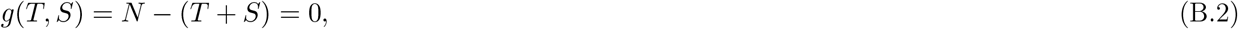

which can then be used in the Lagrangian formulation (B.1), along with our definition of *f*(*S, T*), yielding:

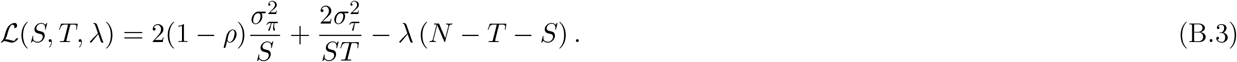

From calculating the partial derivatives of 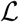 with respect to each variable and setting each derivative to zero, one obtains the following system of equations:

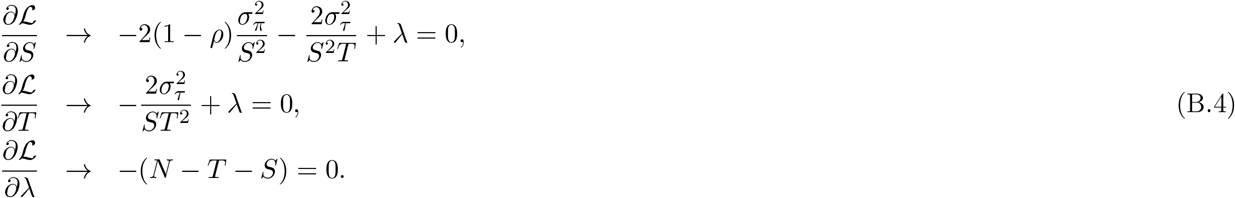

Solving this set of equations (and in the process eliminating λ), one obtains the optimal values of subject and trial sample sizes that minimize *σ*:

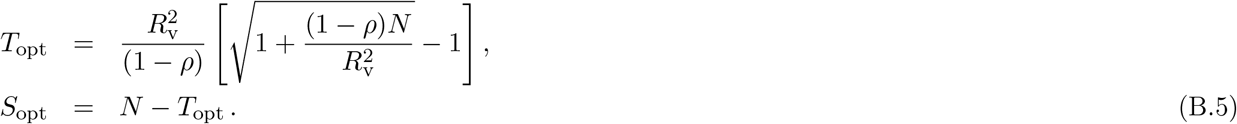

The optimized value of *σ* = *σ*_opt_ can then be obtained by putting these values for *S* and *T* in the variance expression (6). Examples of (*S*_opt_,*T*_opt_) contours are shown in Fig. B.1. In each panel, the same constraint *N* = *T* + *S* is shown with a dashed line, and the optimized (*S*_opt_,*T*_opt_) is shown with a dot, along with the associated isocontour for the optimized *σ*_opt_.

**Figure B.1:**
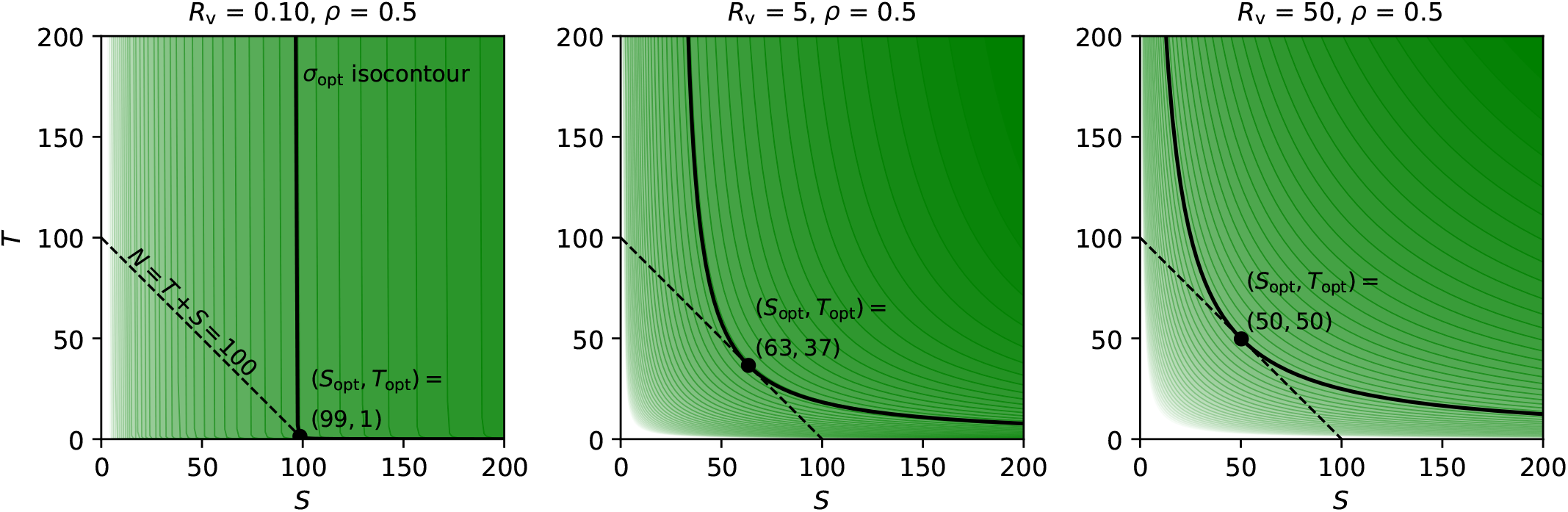
The panels are similar to Fig. 2, showing *σ* isocontours, but here the background opacity increases with increasing statistical efficiency. In each panel, the example constraint *N* = *T* + *S* = 100 is shown with a dashed line, and the optimized (*S*_opt_, *T*_opt_) pair is shown with a dot, along with the associated isocontour for the optimized *σ*_opt_.

**Figure B.2:**
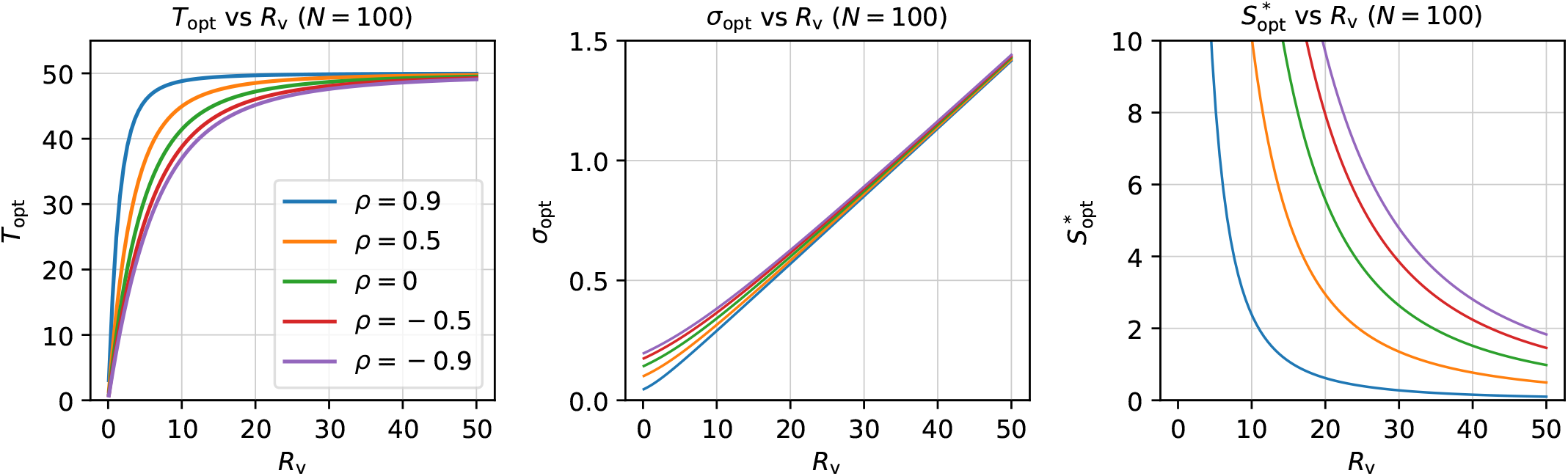
Visualizations of the information in the formula (B.5). Given *N* = 100 total samples, what is the optimal number to partition as trials? The answer depends strongly on the variability ratio *R*_v_: for very low *R*_v_, the optimal number of trials is relatively low; as *R*_v_ increases, *T*_opt_ approaches *N*/2 (where *T*_opt_ = *S*_opt_). The behavior is similar across correlation values, with *ρ* primarily affecting the rate at which *T*_opt_ reaches *N*/2. The middle and right panels show how the optimal uncertainty *σ*_opt_ and minimal subject sample size 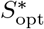 change as functions of *R*_v_ and *ρ*.

Fig. B.2 presents the information in the formula (B.5) in additional ways. In the first panel, the strong relation between *T*_opt_ and *R*_v_ is apparent: for very small variability ratios, the optimal number of trials is near zero; but as *R*_v_ increases, *T*_opt_ → *N*/2, meaning that dividing the samples even between trials and subjects optimizes the statistical uncertainty. The middle panel shows the near linear rise in *σ* with *R*_v_. For small values of *R*_v_, the correlation *ρ* has some impact on the *σ* values, but for larger variability ratios the correlation becomes inconsequential. Finally, the right panel shows the “shift” or “imbalance” term *S*\*, which decreases strongly with *R*_v_.

While the relations in the formula (B.5) are not intuitively obvious, we can understand the analytic behavior in limiting cases, parallel to those in the main text. First, we can investigate the Case 1 limit of small variability ratio *R*_v_, which in this case is compared with the total number of samples *N*. In the specified limit, the following derivations approximate the optimal *T, S* and *σ*:

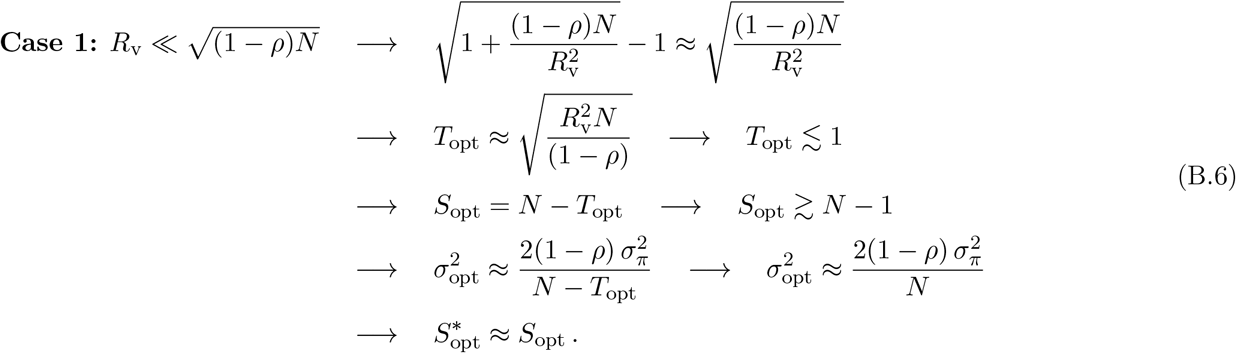

In the above limit, *T*_opt_ shrinks toward zero mathematically, but we have used the notation “≲ 1” to denote that in practice, the number of trial samples cannot become less than 1. As a corollary, *S*_opt_ approaches *N* but in practice would need to be halted at “*N* – 1”. Finally, we note that in this limit 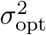 is basically independent of *σ_τ_* and *T*. This case and these features are reflected in the first panel of Fig. B.1.

In the opposite case of large variability ratio, one has the following relations:

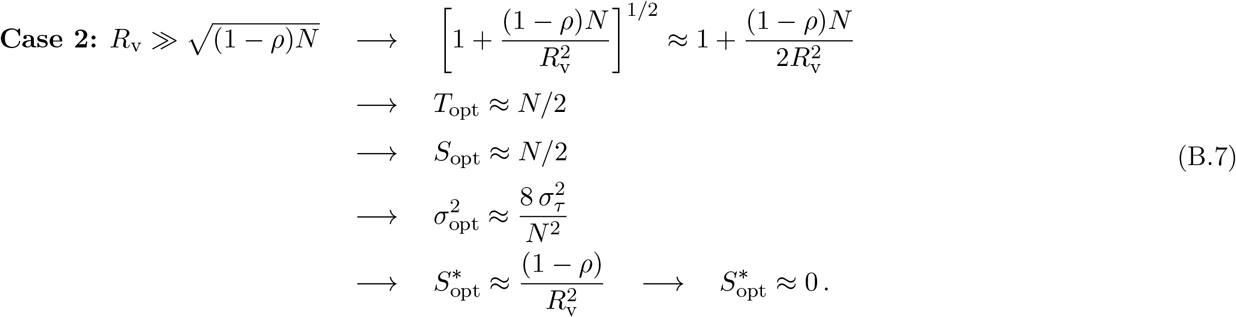

In this limit, the optimized efficiency occurs when the total number of samples is equally divided into trials and samples (as a corollary, the “imbalance” towards subject number *S*\* shrinks toward zero). Then, the optimized efficiency depends only on the cross-trial variability and the product of the two sample sizes. This case and these features are reflected in the last panel of Fig. B.1.

## Footnotes

1 The subsequent discussion would still largely hold in the generic case of heteroscedasticiy (*σ*_*π*1_ ≠ *σ*_*π*2_).

2 While independence is likely an oversimplification, in practice the deviations from this assumption likely would not impact the results discussed here much.

3 The expression (7) maps directly into the general expression of a hyperbola with variables *x* and *y*: *y* = *A* + *B*/(*x* – *C*), where the other parameters are constants that scale or shift the relationship. It could also be viewed in the symmetric hyperbolic form *xy* = *D* through the transformation *y* = *T* and *x* = *S* – *C*, where *C* and *D* are again some parameters. In these formulations, *C* plays the same role as a shift of the curve, which is discussed in the main text as the important parameter *S*\* in (8).

4 Though, on a developmental note: the seeds of this paper originally sprouted *from* some simulations that investigated different modeling behavior as the number of trials was changed. The observation of interesting and unexpected patterns led to an analytic investigation, seeking to understand what was happening on theoretical grounds. Thus, these simulations were developed independently from the analytic understanding and not created simply to prove the preceding equations. And, for those of us who are typically theoreticians, it has also been good to have a reminder of the power of experimental and simulation work.

5 As evaluated using the AFNI GUI’s “whereami” functionality, referencing the Glasser MNI atlas (Glasser et al., 2016)

## References

Cai, C., Sekihara, K., Nagarajan, S.S., 2018. Hierarchical multiscale Bayesian algorithm for robust MEG/EEG source reconstruction. NeuroImage 183, 698–715.

Chen, G., Padmala, S., Chen, Y., Taylor, P.A., Cox, R.W., Pessoa, L., 2020. To pool or not to pool: Can we ignore cross-trial variability in FMRI? NeuroImage, 117496.

Chen, G., Pine, D.S., Brotman, M.A., Smith, A.R., Cox, R.W., Haller, S.P., 2021. Trial and error: A hierarchical modeling approach to test-retest reliability. NeuroImage 245, 118647.

Chen, G., Saad, Z.S., Britton, J.C., Pine, D.S., Cox, R.W., 2013. Linear mixed-effects modeling approach to FMRI group analysis. NeuroImage 73, 176–190.

Chen, G., Saad, Z.S., Nath, A.R., Beauchamp, M.S., Cox, R.W., 2012. FMRI group analysis combining effect estimates and their variances. NeuroImage 60, 747–765.

Chiang, S., Guindani, M., Yeh, H.J., Dewar, S., Haneef, Z., Stern, J.M., Vannucci, M., 2017. A Hierarchical Bayesian Model for the Identification of PET Markers Associated to the Prediction of Surgical Outcome after Anterior Temporal Lobe Resection. Frontiers in Neuroscience 11, 669.

Cox, R.W., 1996. AFNI: Software for analysis and visualization of functional magnetic resonance neuroimages. Comput Biomed Res 29, 162–173.

Davis, T., LaRocque, K.F., Mumford, J.A., Norman, K.A., Wagner, A.D., Poldrack, R.A., 2014. What do differences between multi-voxel and univariate analysis mean? how subject-, voxel-, and trial-level variance impact fMRI analysis. NeuroImage 97, 271–283.

Desmond, J.E., Glover, G.H., 2002. Estimating sample size in functional MRI (fMRI) neuroimaging studies: Statistical power analyses. Journal of Neuroscience Methods 118, 115–128.

Durnez, J., Degryse, J., Moerkerke, B., Seurinck, R., Sochat, V., Poldrack, R.A., Nichols, T.E., 2016. Power and sample size calculations for fMRI studies based on the prevalence of active peaks. bioRxiv, 049429.

Eriksen, B.A., Eriksen, C.W., 1974. Effects of noise letters upon the identification of a target letter in a nonsearch task. Perception & Psychophysics 16, 143–149.

Glasser, M.F., Coalson, T.S., Robinson, E.C., Hacker, C.D., Harwell, J., Yacoub, E., Ugurbil, K., Andersson, J., Beckmann, C.F., Jenkinson, M., Smith, S.M., Van Essen, D.C., 2016. A multi-modal parcellation of human cerebral cortex. Nature 536, 171–178.

Gonzalez-Castillo, J., Saad, Z.S., Handwerker, D.A., Inati, S.J., Brenowitz, N., Bandettini, P.A., 2012. Wholebrain, time-locked activation with simple tasks revealed using massive averaging and model-free analysis. PNAS 109, 5487–5492.

Gordon, E.M., Laumann, T.O., Gilmore, A.W., Newbold, D.J., Greene, D.J., Berg, J.J., Ortega, M., Hoyt-Drazen, C., Gratton, C., Sun, H., Hampton, J.M., Coalson, R.S., Nguyen, A.L., McDermott, K.B., Shimony, J.S., Snyder, A.Z., Schlaggar, B.L., Petersen, S.E., Nelson, S.M., Dosenbach, N.U.F., 2017. Precision Functional Mapping of Individual Human Brains. Neuron 95, 791–807.e7.

Haines, N., Kvam, P.D., Irving, L.H., Smith, C., Beauchaine, T.P., Pitt, M.A., Ahn, W.Y., Turner, B., 2020. Learning from the Reliability Paradox: How Theoretically Informed Generative Models Can Advance the Social, Behavioral, and Brain Sciences. Preprint. PsyArXiv. doi:10.31234/osf.io/xr7y3.

He, B.J., Zempel, J.M., 2013. Average Is Optimal: An Inverted-U Relationship between Trial-to-Trial Brain Activity and Behavioral Performance. PLOS Computational Biology 9, e1003348.

Molloy, M.F., Bahg, G., Li, X., Steyvers, M., Lu, Z.L., Turner, B.M., 2018. Hierarchical Bayesian Analyses for Modeling BOLD Time Series Data. Comput Brain Behav 1, 184–213.

Mumford, J.A., 2012. A power calculation guide for fMRI studies. Social Cognitive and Affective Neuroscience 7, 738–742.

Naselaris, T., Allen, E., Kay, K., 2021. Extensive sampling for complete models of individual brains. Current Opinion in Behavioral Sciences 40, 45–51.

Nee, D.E., 2019. fMRI replicability depends upon sufficient individual-level data. Commun Biol 2, 1–4.

Ostwald, D., Schneider, S., Bruckner, R., Horvath, L., 2019. Power, positive predictive value, and sample size calculations for random field theory-based fMRI inference. bioRxiv, 613331.

Rohe, T., Ehlis, A.C., Noppeney, U., 2019. The neural dynamics of hierarchical Bayesian causal inference in multisensory perception. Nat Commun 10, 1907.

Rouder, J., Kumar, A., Haaf, J.M., 2019. Why Most Studies of Individual Differences With Inhibition Tasks Are Bound To Fail. Technical Report. PsyArXiv. doi:10.31234/osf.io/3cjr5.

Rouder, J.N., Haaf, J.M., 2019. A psychometrics of individual differences in experimental tasks. Psychon Bull Rev 26, 452–467.

Smith, A.R., White, L.K., Leibenluft, E., McGlade, A.L., Heckelman, A.C., Haller, S.P., Buzzell, G.A., Fox, N.A., Pine, D.S., 2020. The Heterogeneity of Anxious Phenotypes: Neural Responses to Errors in Treatment-Seeking Anxious and Behaviorally Inhibited Youths. Journal of the American Academy of Child & Adolescent Psychiatry 59, 759–769.

Szucs, D., Ioannidis, J.P., 2020. Sample size evolution in neuroimaging research: An evaluation of highly-cited studies (1990–2012) and of latest practices (2017–2018) in high-impact journals. NeuroImage 221, 117164.

Thirion, B., Pinel, P., Mériaux, S., Roche, A., Dehaene, S., Poline, J.B., 2007. Analysis of a large fMRI cohort: Statistical and methodological issues for group analyses. NeuroImage 35, 105–120.

Trenado, C., González-Ramírez, A., Lizárraga-Cortés, V., Pedroarena Leal, N., Manjarrez, E., Ruge, D., 2019. The Potential of Trial-by-Trial Variabilities of Ongoing-EEG, Evoked Potentials, Event Related Potentials and fMRI as Diagnostic Markers for Neuropsychiatric Disorders. Front. Neurosci. 12.

Turner, B.M., Palestro, J.J., Miletić, S., Forstmann, B.U., 2019. Advances in techniques for imposing reciprocity in brain-behavior relations. Neuroscience & Biobehavioral Reviews 102, 327–336.

Turner, B.M., Rodriguez, C.A., Norcia, T.M., McClure, S.M., Steyvers, M., 2016. Why more is better: Simultaneous modeling of EEG, fMRI, and behavioral data. NeuroImage 128, 96–115.

Turner, B.O., Paul, E.J., Miller, M.B., Barbey, A.K., 2018. Small sample sizes reduce the replicability of task-based fMRI studies. Commun Biol 1, 1–10.

Webb-Vargas, Y., Chen, S., Fisher, A., Mejia, A., Xu, Y., Crainiceanu, C., Caffo, B., Lindquist, M.A., 2017. Big Data and Neuroimaging. Stat Biosci 9, 543–558.

Westfall, J., Kenny, D.A., Judd, C.M., 2014. Statistical power and optimal design in experiments in which samples of participants respond to samples of stimuli. J Exp Psychol Gen 143, 2020–2045.

Westfall, J., Nichols, T.E., Yarkoni, T., 2017. Fixing the stimulus-as-fixed-effect fallacy in task fMRI. Wellcome Open Res 1.

Wolff, A., Chen, L., Tumati, S., Golesorkhi, M., Gomez-Pilar, J., Hu, J., Jiang, S., Mao, Y., Longtin, A., Northoff, G., 2021. Prestimulus dynamics blend with the stimulus in neural variability quenching. NeuroImage 238, 118160.

